# Comprehensive structure-function characterization of DNMT3B and DNMT3A reveals distinctive *de novo* DNA methylation mechanisms

**DOI:** 10.1101/2020.04.27.064386

**Authors:** Linfeng Gao, Max Emperle, Yiran Guo, Sara A Grimm, Wendan Ren, Sabrina Adam, Hidetaka Uryu, Zhi-Min Zhang, Dongliang Chen, Jiekai Yin, Michael Dukatz, Hiwot Anteneh, Renata Z. Jurkowska, Jiuwei Lu, Yinsheng Wang, Pavel Bashtrykov, Paul A Wade, Gang Greg Wang, Albert Jeltsch, Jikui Song

**Author notes:** These authors contributed equally to this work (Linfeng Gao, Max Emperle, Yiran Guo). School of pharmacy, Jinan University, 601 Huangpu Avenue West, Guangzhou 510632, China (Zhi-Min Zhang). School of Biosciences, Cardiff University, Sir Martin Evans Building, Museum Avenue, Cardiff, CF10 3AX, UK.

## Abstract

Mammalian DNA methylation patterns are established by two *de novo* DNA methyltransferases DNMT3A and DNMT3B, which exhibit both redundant and distinctive methylation activities. However, the related molecular basis remains undetermined. Through comprehensive structural, enzymology and cellular characterization of DNMT3A and DNMT3B, we here report a multi-layered substrate-recognition mechanism underpinning their divergent genomic methylation activities. A hydrogen bond in the catalytic loop of DNMT3B causes a lower CpG specificity than DNMT3A, while the interplay of target recognition domain and homodimeric interface fine-tunes the distinct target selection between the two enzymes, with Lysine 777 of DNMT3B acting as a unique sensor of the +1 flanking base. The divergent substrate preference between DNMT3A and DNMT3B provides an explanation for site-specific epigenomic alterations seen in ICF syndrome with DNMT3B mutations. Together, this study reveals crucial and distinctive substrate-readout mechanisms of the two DNMT3 enzymes, implicative of their differential roles during development and pathogenesis.

## Introduction

DNA methylation is one of the major epigenetic mechanisms that critically influence gene expression, genomic stability and cell differentiation^1-3^. In mammals, DNA methylation predominantly occurs at the C-5 position of cytosine within the symmetric CpG dinucleotide, affecting ∼70-80% of the CpG sites throughout the genome4. Mammalian DNA methylation patterns are mainly generated by two *de novo* DNA methyltransferases, DNMT3A and DNMT3B^5^. The catalytically inactive DNMT3-like protein (DNMT3L) has an important regulatory role in this process by acting as cofactor of DNMT3A or DNMT3B^6-8^. In addition to CpG methylation, DNMT3A and DNMT3B introduce non-CpG methylation (mainly CpA) in oocytes, embryonic stem (ES) cells and neural cells^4, 9-11^. The presence of non-CpG methylation was reported to correlate with transcriptional repression^12, 13^ and the pluripotency-associated epigenetic state^4, 14^, lending support for non-CpG methylation as an emerging epigenetic mark in defining tissue-specific patterns of gene expression, particularly in the brain.

DNMT3A and DNMT3B are closely related in amino acid sequence^5^, with a C-terminal methyltransferase (MTase) domain preceded by regulatory regions including a proline-tryptophan-tryptophan-proline (PWWP) domain and an ATRX-DNMT3-DNMT3L-type (ADD) zinc finger domain^15, 16^. Previous studies have indicated a partial redundancy between the two enzymes in the establishment of methylation patterns across the genome^5, 17^; however, a single knockout (KO) of either DNMT3A or DNMT3B resulted in embryonic or postnatal lethality, indicating their functional distinctions^5, 17-19^. Indeed, it was shown that DNMT3A is critical for establishing methylation at major satellite repeats and allele-specific imprinting during gametogenesis^8, 17^, whereas DNMT3B plays a dominant role in early embryonic development and in minor satellite repeat methylation^5, 17^. Mutations of DNMT3A are prevalent in hematological cancers such as acute myeloid leukemia (AML) ^20^ and occur in a developmental overgrowth syndrome^21^; in contrast, mutations of DNMT3B lead to the Immunodeficiency, centromeric instability, facial anomalies (ICF) syndrome^5, 22, 23, 24^. Previous studies have indicated subtle mechanistic differences between DNMT3A and DNMT3B^9, 25-28^, including their differential preference toward the flanking sequence of CpG target sites^29-32^. However, due to the limited number of different substrates investigated in these studies, global differences in substrate recognition of DNMT3A and DNMT3B remain elusive. Our recently reported crystal structure of the DNMT3A-DNMT3L heterotetramer in complex with CpG DNA^33^ revealed that the two central DNMT3A subunits bind to the same DNA duplex through a set of interactions mediated by protein motifs from the target recognition domain (TRD), the catalytic core and DNMT3A-DNMT3A homodimeric interface (also called RD interface below). However, the structural basis of DNMT3B-mediated methylation remains unclear.

To gain mechanistic understanding of *de novo* DNA methylation, we here report comprehensive enzymology, structural and cellular characterizations of DNMT3A- and DNMT3B-complexes. Our results uncover their distinct substrate and flanking sequence preferences, implicating epigenomic alterations caused by DNMT3 mutations in diseases. Notably, we show that the catalytic core, TRD domain and RD interface cooperate in orchestrating a distinct, multi-layered substrate-readout mechanism between DNMT3A and DNMT3B, which impacts the establishment of CpG and non-CpG methylation patterns in cells.

## Results

### Deep enzymology analysis of DNMT3A and DNMT3B

To systematically elucidate the functional divergence of DNMT3A and DNMT3B, we have developed a deep enzymology workflow to study the substrate specificity of DNA MTases in random sequence context. In essence, we generated a pool of DNA substrates, in which the target CpG site is flanked by 10 random nucleotides on each side. Following methylation by the MTase domains of DNMT3A or DNMT3B, the reaction products were subjected to hairpin ligation, bisulfite conversion, PCR amplification and next generation sequencing (NGS) analysis (Supplementary Fig. 1 and Supplementary Table 1). Analysis of the base enrichments at all flank positions in the methylated sequences demonstrated that DNMT3A- and DNMT3B-mediated methylation is significantly influenced by the CpG-flanking sequence from the −2 to the +3 site (Fig. 1a). Based on this result, we focused on further analyzing the effect of the ±3 bp flanking positions on the activity of both enzymes. Methylation levels were averaged for all 4096 NNNCGNNN sites for the two experiments with DNMT3A and DNMT3B, revealing high bisulfite conversion (>99.5%) (Supplementary Table 1), high coverage of NNNCGNNN sites that was similar between DNMT3A and DNMT3B (Supplementary Fig. 2a-d) and a high correlation of average methylation levels for the different flanks between experimental repeats, yet different between DNMT3A and DNMT3B (Fig. 1b and Supplementary Fig. 2e). We then validated these observations by *in vitro* methylation analysis on 30-mer oligonucleotide substrates^34^ with CpG sites in different trinucleotide flanking sequence context, which reveals methylation preferences of both enzymes in strong agreement with the results from the NGS profile (Supplementary Fig. 3a, b).

**Figure 1.**
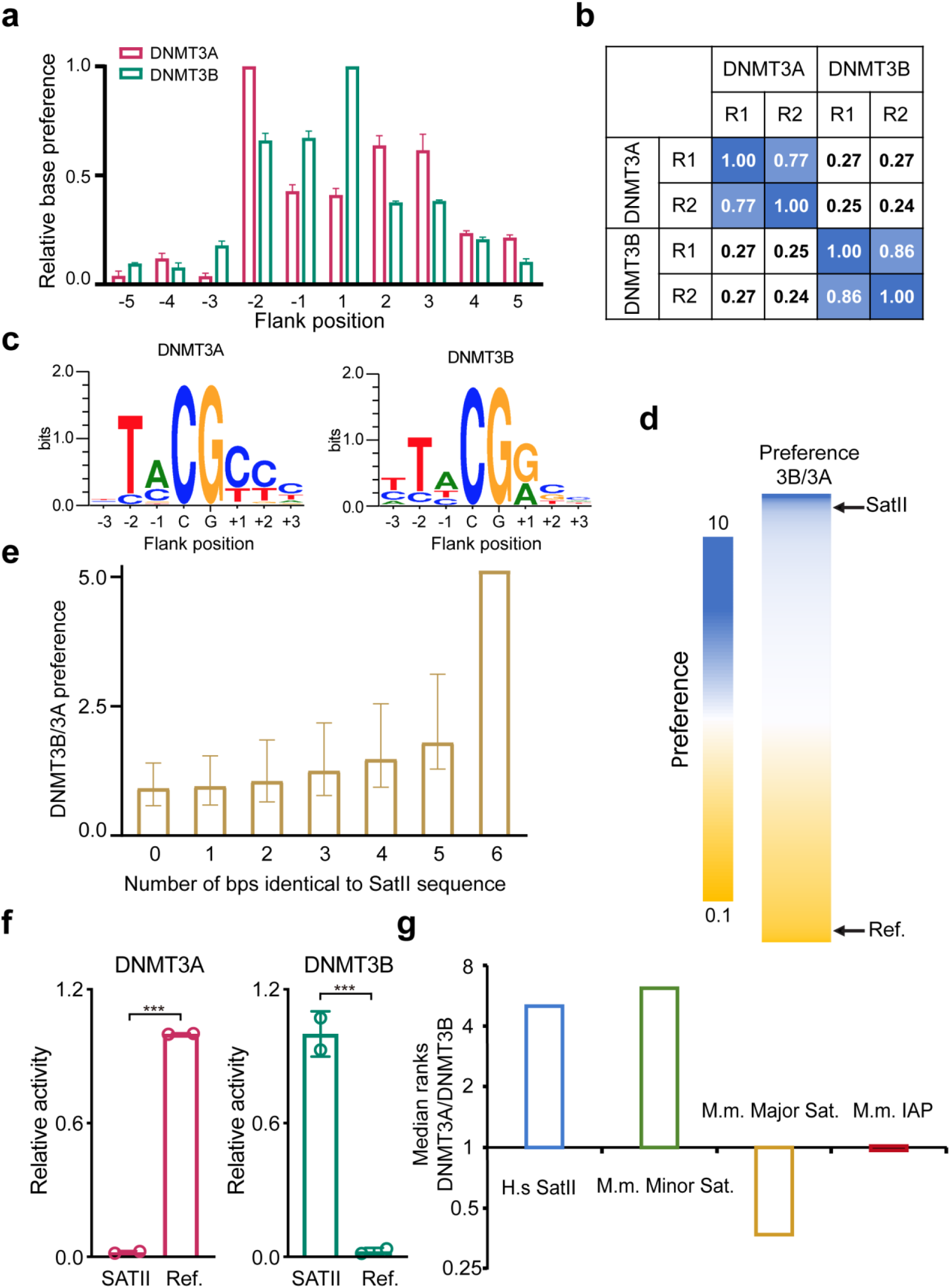
Deep enzymology flanking sequence analysis links intrinsic substrate preference of DNMT3B to SatII sequence recognition. **(a)** Relative base preferences at the −5 to +5 flanking positions of mouse DNMT3A (mDNMT3A) and DNMT3B (mDNMT3B) indicating the strength of sequence readout at each site. The numbers refer to the standard deviations of the observed/expected base composition at each site among the methylated sequence reads, normalized to the highest value for each enzyme. **(b)** Correlation of the NNNCGNNN methylation profiles of mDNMT3A and mDNMT3B for two independent experiments (repetition R1 and R2). The numbers refer to the pairwise Pearson correlation coefficients. **(c)** Weblogos of the 50-200 most preferred NNNCGNNN methylation sites by mDNMT3A and mDNMT3B. Quantitative enrichment and depletion plots and statistics are provided in Supplementary Fig. 2c, d. **(d)** Heatmap of the normalized DNMT3B/DNMT3A preferences for methylation of NNNCGNNN sites. The position of the SatII sequence (TCCATT<u>CG</u>ATGATG) in the ranking is indicated (rank 131 of 4096) as well as the position of the mDNMT3B-disfavored reference substrate (AGG<u>CG</u>CCC) used in panel C (rank 4092 of 4096). **(e)** mDNMT3B/mDNMT3A preferences (B/A preference) were binned and averaged based on their similarity to the SatII sequence. The median preference of each bin is shown. The error bars indicate the first and third quartile. **(f)** Experimental validation of the SatII preference of DNMT3B by radioactive methylation assays. The methylation activity of each enzyme was normalized to the more active substrate. The figure shows average values and standard deviations based on three independent measurements (see also Supplementary Fig. S6a, b). Statistical analysis used two-tailed Student’s t-test for the difference of activities: ***, *p* < 0.001. **(g)** Ratio of the medians of the distributions of mDNMT3A/mDNMT3B preference among all CpG sites from mouse minor satellite repeats (Genbank Z22168.1), major satellite repeats (Genbank EF028077.1) and IAP elements (Genbank AF303453.1). A high DNMT3A/DNMT3B rank ratio corresponds to preferred methylation by DNMT3B. Refer to Supplementary Fig. 6d and the text for details.

Strikingly, the effect of flanking sequences on methylation rates of DNMT3A and DNMT3B is very pronounced showing NNNCGNNN flanks with very high and very low methylation in both data sets (Supplementary Table 2). Weblogo analysis of the most highly methylated sites revealed that DNMT3A shows a preference toward a CG(C/T) motif, whereas DNMT3B shows a higher activity toward a CG(G/A) motif (Fig. 1c). In addition, both enzymes prefer a T at the −2 flank site. To explore the differences between DNMT3A and DNMT3B systematically, we computed the ratio of the sequence preferences of DNMT3B and DNMT3A (Fig. 1d and Supplementary Table 3) designated as B/A preference from here on. Overall, these preference ratios span a more than 100-fold range, illustrating the large divergence in flanking sequence preferences between the two enzymes. These data provide for the first time a systematic survey of the substrate preferences of DNMT3A and DNMT3B and highlight their distinct enzymatic properties.

### Genomic profiling of cellular methylation introduced by DNMT3B and DNMT3A

To examine patterns of *de novo* DNA methylation induced by DNMT3A or DNMT3B in cells, we stably transduced either enzyme into mouse ES cells with compound knockout (TKO) of DNMT1, DNMT3A and DNMT3B as carried out before (Supplementary Fig. 4a)^33^. Using liquid chromatography–mass spectrometry (LC-MS)-based quantification, we detected a global increase in cytosine methylation in TKO cells after rescue with DNMT3B (Supplementary Fig. 4b), an effect similar to what was observed with DNMT3A rescue^33^. Next, we profiled the genome-wide methylation introduced by either DNMT3A or DNMT3B in cells by enhanced reduced representation bisulfite sequencing (eRRBS, two replicates per group; Supplementary Table 4) and generated datasets with desired high conversion rates (Supplementary Fig. 4c) and reproducibility between replicates (Supplementary Fig. 4d). Next, we aimed to compare deep enzymology and eRRBS profiling results. Here, average eRRBS methylation levels within the NNNCGNNN sequence contexts were computed from either DNMT3A- or DNMT3B-reconstituted cells, and the averaged eRRBS methylation levels in both samples were found to be high and equal in coverage (Supplementary Fig. 5a, b). The eRRBS methylation levels were then compared with the biochemical activities of DNMT3B and DNMT3A (Supplementary Fig. 5c). We found that the biochemical activity of DNMT3B correlated positively with the genomic methylation of DNMT3B, but not with the genomic methylation of DNMT3A; conversely, the biochemical activity of DNMT3A correlated well with the genomic methylation of DNMT3A, but not with that of DNMT3B (Supplementary Fig. 5c and Supplementary Table 3). Hence, despite being different assay systems, eRRBS and *in vitro* methylation based analyses yielded consistent results, highlighting a strong effect of the flanking sequence on DNMT3B and DNMT3A activities in cells.

### Methylation of repetitive sequences by DNMT3A and DNMT3B

We have further interrogated the biochemical B/A substrate preference with one of the best-characterized DNMT3B targets, the SatII repeats, methylation of which is lost in the ICF syndrome^24^. Toward this, we used the ATTCGATG consensus sequence of SatII repeats^35^ for analysis. Notably, the ATTCGATG sequence is ranked 131^th^ among the 4096 possible NNNGCNNN sequences in the biochemical B/A preferences, where a low rank corresponds to high preference (Fig. 1d). Next, by averaging the B/A preferences for the NNNCGNNN sequences, grouped by their similarity to the SatII sequence, we observed a strong correlation of the averaged B/A preferences with an increasing similarity to the SatII sequence, illustrating the adaptation of DNMT3B for SatII methylation (Fig. 1e). Furthermore, we assayed human DNMT3A- and DNMT3B-mediated methylation on 30-mer oligonucleotide substrates^34^ containing a CpG with either the SatII 6-bp flanking sequence (TCCATTCGATGATG) or a DNMT3B-disfavored reference substrate (AGGCGCCC, rank 4092 of 4096 in the B/A preference, Fig.1d). Strikingly, DNMT3B showed an ∼11-fold higher activity on the SatII substrate, whereas DNMT3A was ∼15-fold more active on the reference substrate (Fig. 1f and Supplementary Fig. 6a, b). Similar enzymatic preferences were observed for mouse DNMT3A and DNMT3B (Supplementary Fig. 6c). These data confirm the specific activity of DNMT3B on the SatII targets and document a >100 difference in methylation rates of substrates with different flanking sequences by DNMT3B and DNMT3A.

We then analyzed the sequences of murine genomic repeats to determine the ranks of CpG-associated sequences in the sorted preferences of DNMT3A and DNMT3B. For this only the more preferred DNA strand was considered, because DNMT1-mediated maintenance DNA methylation in cells would rapidly methylate hemimethylated CpG sites regardless of which DNA strand was initially methylated. This analysis reproduced the preference of DNMT3B for human SatII sequences, and remarkably mirrored genomic methylation data of DNMT3A and DNMT3B in mice^17^ where a pronounced preference of DNMT3B and DNMT3A was observed for minor satellite repeats and major satellite repeats, respectively, whereas both enzymes showed equal preferences for the IAP sequence (Fig. 1g and Supplementary Fig. 6d).

### DNMT3A- and DNMT3B-mediated non-CpG methylation

Our deep enzymology and eRRBS studies have further allowed direct comparison between the DNMT3A- and DNMT3B-mediated CpG and CpH methylation. Intriguingly, analysis of the CpA/CpG ratio in the NNNCGNNN context for each enzyme reveals that DNMT3A CpG specificity is higher within a favored flanking sequence context, whereas in the case of DNMT3B, a preferable CpG flanking context is correlated with a decreased CpG specificity and higher CpA methylation (Supplementary Fig. 6e,f), which is in line with the stronger correlation of flanking sequence profiles of CpG and CpA methylation for DNMT3B than for DNMT3A (Supplementary Fig. 6g). A targeted activity analysis revealed that the preference of DNMT3B for a G at the +1 site is strongly enhanced for non-CpG methylation, but not that of DNMT3A (Fig. 2a,b). Furthermore, our eRRBS-based methylome data (Supplementary Fig. 7) obtained from DNMT3A or DNMT3B-reconstituted TKO cells also revealed most efficient CpH methylation of sequences containing C and G on the +1 site, respectively (Fig. 2c,d) in agreement with the activity data and previously published cellular methylation data^4, 36-40^. Together, these data provide a direct link between the intrinsic substrate specificities of DNMT3s and the DNA methylation patterns in cells.

**Figure 2.**
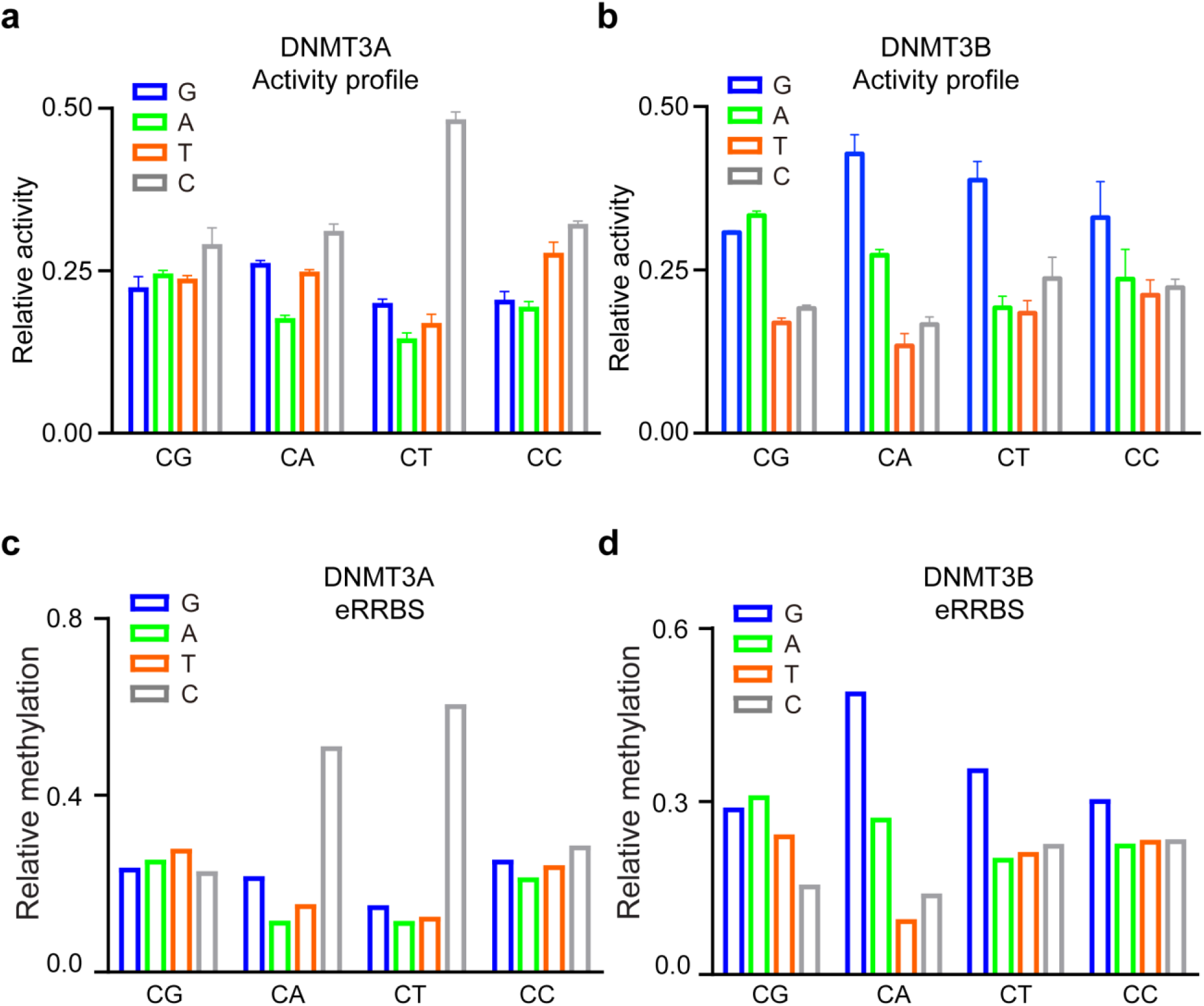
Non-CpG methylation by DNMT3A and DNMT3B. **(a**,**b)** CpG and non-CpG methylation of mDNMT3A (**a**) and mDNMT3B (**b**) averaged for the different +1 flanking base pairs. The cytosine-containing dinucleotides are displayed on the horizontal axis and +1 base is shown in the different bars, respectively. Error bars indicate the SD of two experimental repeats. **(c**,**d)** eRRBS analysis of CpG and non-CpG methylation in TKO cells introduced by hDNMT3A (**c**) and hDNMT3B (**d**) averaged for the different +1 flanking base pairs (depth of reads >10 and *p* < 0.0001). The cytosine-containing dinucleotides are displayed on the horizontal axis and +1 base is shown in the different bars.

### Crystal structures of the DNMT3B-DNMT3L-CpG DNA complexes

To gain a mechanistic understanding of the enzymatic divergence of DNMT3A and DNMT3B, we generated the complexes of DNMT3B-DNMT3L, formed by the MTase domain of human DNMT3B and the C-terminal domain of human DNMT3L, with two different Zebularine(Z)-containing DNA duplexes harboring two ZpGpA or ZpGpT sites as mimics of CGA and CGT motifs, which are separated by 14 bps (Fig. 3a and Supplementary Fig. 8a-c). The structures of the DNMT3B-DNMT3L-CGA DNA (DNMT3B-CGA) and DNMT3B-DNMT3L-CGT DNA (DNMT3B-CGT) bound to the cofactor product *S*-Adenosyl-L-homocysteine (SAH) were both solved at 3.0 Å resolution (Table 1). The two structures are very similar, with a root-mean-square deviation (RMSD) of 0.27 Å over 828 Cα atoms (Supplementary Fig. 8d). For simplification, we choose the DNMT3B-CGA complex for structural analysis, unless indicated otherwise. Similar to the DNMT3A-DNMT3L complexes^33, 41, 42^, the DNMT3B-DNMT3L-DNA complex reveals a linear heterotetrameric arrangement of DNMT3L-DNMT3B-DNMT3B-DNMT3L (Fig. 3b).

**Figure 3.**
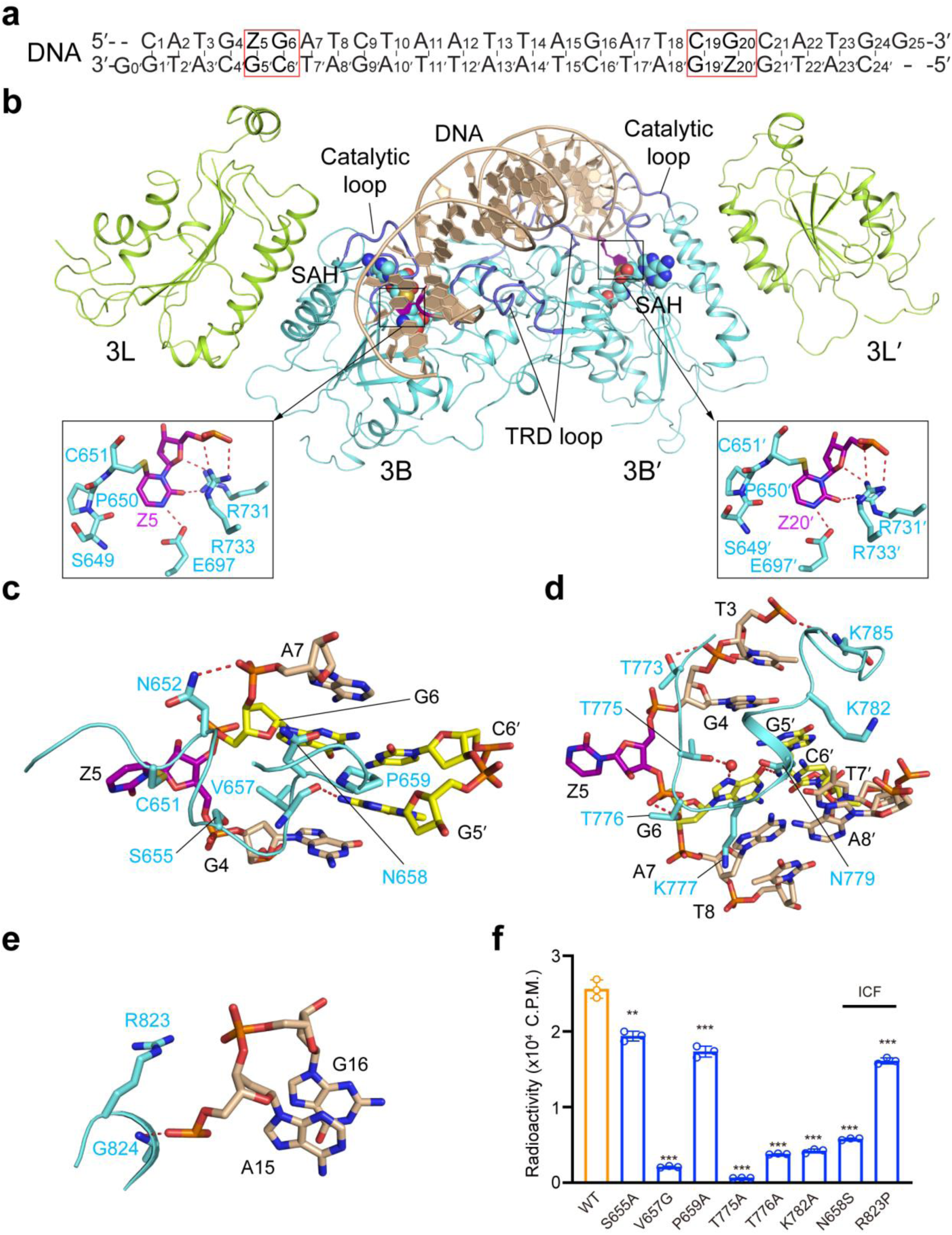
Structure of the DNMT3B-DNMT3L tetramer in complex with CGA DNA. **(a)** DNA sequence (CGA) used for the structural study. Z, Zebularine, a cytidine analog that is known to form stable covalent complexes with DNMTs^46^. **(b)** Ribbon representations of human DNMT3B-DNMT3L bound to DNA and SAH. The DNA-contacting TRD loop and catalytic loop are colored in slate. The Zebularines anchored at the two active sites are shown in expanded views, with hydrogen-bonding interactions depicted as dashed lines. For identification of hydrogen bonds, the upper limit of the donor-acceptor distance is set to 3.5 Å. **(c-e)** Close-up views of the DNA interactions of the catalytic loop (**b**), TRD loop (**c**) and the loop at the RD interface (**d**) of DNMT3B. The DNA is colored in wheat, except for the ZpG site, in which the Zebularine and the other three nucleotides are colored in purple and yellow, respectively. The hydrogen bonding interactions are shown as dashed lines. **(f)** *In vitro* methylation of wild-type (WT) and mutant DNMT3B in the form of DNMT3B-DNMT3L tetramer analyzed by radioactive methylation assays. Data are mean ± SD. Statistical analysis used two-tailed Student’s t-test for the difference from WT: **, *p* < 0.01; ***, *p* < 0.001.

The interaction between DNMT3B and DNA involves a loop from the catalytic core (catalytic loop: residues 648-672), a loop from the TRD (TRD loop: residues 772-791), and a segment in the homodimeric interface of DNMT3B, known as the RD interface in DNMT3A^42^ (RD: residues 822-828) (Fig. 3c-e and Supplementary Fig. 8e,f). Each Zebularine (Z5/Z20′) is flipped into the active site of one DNMT3B subunit, where it is anchored through a covalent linkage with the catalytic cysteine C651 and hydrogen-bonding interactions with other catalytic residues (Fig. 3b). The cavities vacated by the base flipping of Z5/Z20′ are occupied by the side chains of V657 (Fig. 3c). The orphan guanine (G5′/G20) that originally paired with Z5/Z20′ is stabilized by a hydrogen bond between the backbone carbonyl group of V657 and the Gua-N2 atom (Fig. 3c), while the ZpG guanine (G6/G19′) is recognized by a hydrogen bond between the side chain of N779 and the Gua-O6 atom, as well as a water-mediated hydrogen bond between the side chain of T775 and the Gua-N7 atom (Fig. 3d). Both guanines also engage van der Waals contacts with catalytic loop residue P659 (Fig. 3c). Aside from the CpG recognition, N779 forms a hydrogen bond with the T7′-O4 atom (Fig. 3d), suggestive of a role in recognizing the nucleotides at the +1 flank position. Furthermore, residues on the catalytic loop (N652 and S655), the TRD loop (Q772, T773, T776, K782 and K785) and the RD interface (R823 and G824) interact with the DNA backbone on both strands through hydrogen-bonding or electrostatic interactions (Fig. 3c-e and Supplementary Fig. 8e,f). Note that DNMT3L does not make any contact with DNA (Fig. 3b), as previously observed for the DNMT3A-DNMT3L-DNA complex^33^ and it did not show strong effects on the intrinsic flanking sequence preferences of DNMT3A or DNMT3B (Supplementary Fig. 9).

Guided by the structural analysis, we selected a number of DNA-interacting residues of DNMT3B for mutagenesis and enzymatic assays. In comparison with wild-type (WT) DNMT3B, mutation of the catalytic-loop residues (S655A, V657G, N658S and P659A), RD residue (R823P) and the TRD-loop residues (T775A, T776A and K782A) led to a modest to severe decrease in the enzymatic activity (Fig. 3f), particularly for V657G, T776A and K782A, supporting the notion that these residues are important for DNMT3B-substrate recognition and catalysis. Note that two of these mutations, N658S and R823P, are associated with ICF syndrome^26, 43^, which reinforces the etiologic link between DNMT3B and the ICF syndrome.

### Role of the catalytic loop in defining the CpG specificities of DNMT3A and DNMT3B

Structural alignments of the DNMT3B-CGA and DNMT3B-CGT complexes with the DNMT3A-CGT complex (PDB 5YX2) give an RMSD of 0.62 Å and 0.63 Å over 853 and 863 aligned Cα atoms, respectively (Supplementary Fig. 10a), in line with the ∼80% sequence identity between the MTase domains of DNMT3A and DNMT3B (Supplementary Fig. 10b). Nevertheless, a notable conformational difference is observed for the catalytic loop (Fig. 4a): A side-chain hydrogen bond is formed between DNMT3B N656 and R661 (Fig. 4b), but not between the corresponding DNMT3A residues I715 and R720 (Fig. 4c). Such a conformational difference presumably leads to altered dynamic behavior of the catalytic loop, as demonstrated by the subtle conformational repositioning of the CpG-contacting residues in DNMT3B, such as V657 and P659, relative to their DNMT3A counterparts (Fig. 4a).

**Figure 4.**
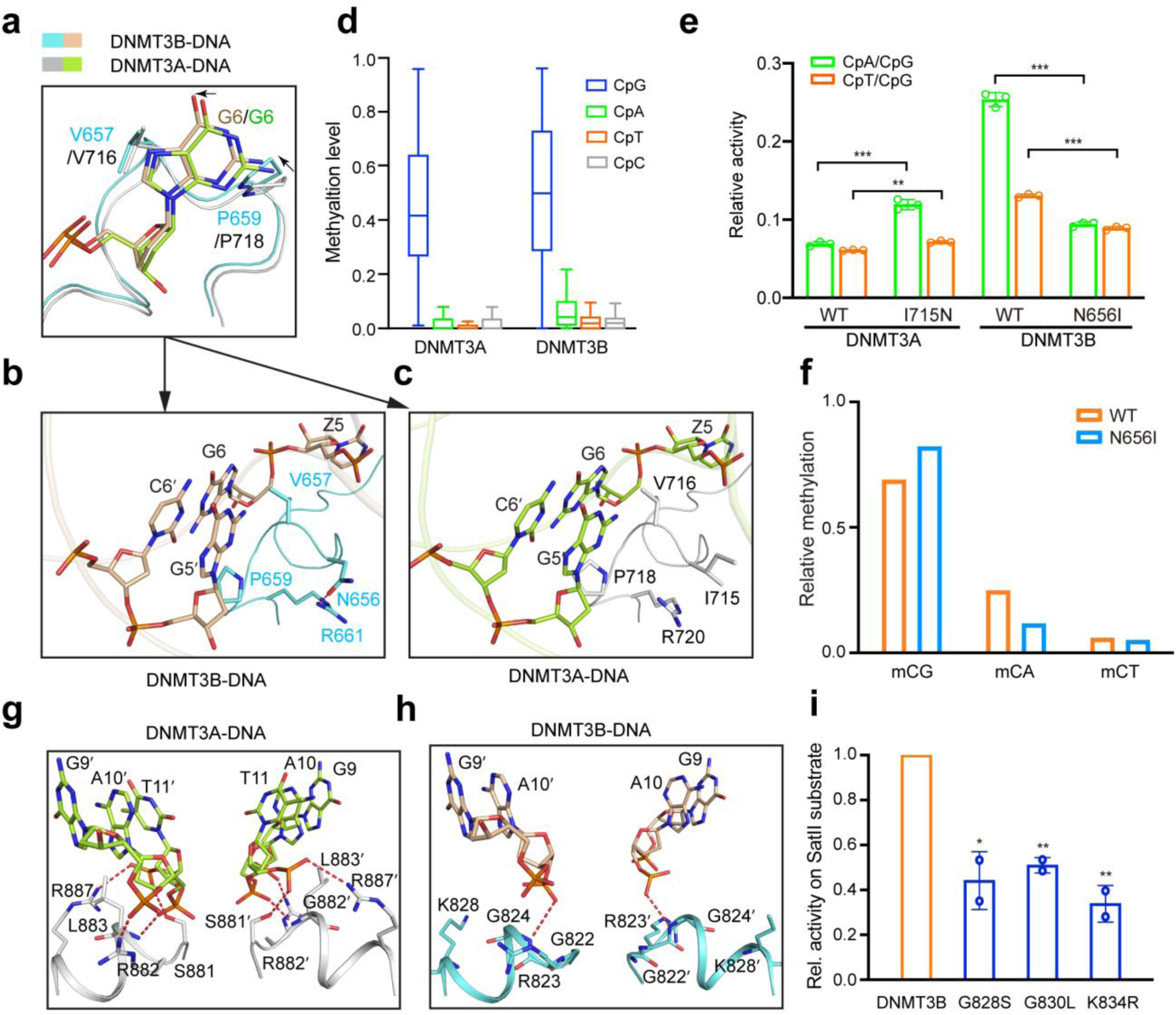
Divergent DNA recognition between DNMT3A and DNMT3B. **(a-c)** Structural comparison of the DNMT3A-DNA and DNMT3B-DNA complexes, highlighting distinct intramolecular interactions within the catalytic loops of DNMT3A (**b**) and DNMT3B (**c**). **(d)** Box plot of CpG and non-CpG methylation in random flanking context. The boxes display the 1^st^ and 3^rd^ quartiles with medians indicated. Whiskers display the data range. **(e)** *In vitro* CpG and CpH methylation of hDNMT3A-hDNMT3L and hDNMT3B-hDNMT3L, WT or mutants on the catalytic loop, using (GAC)_12_, (AAC)_12_ and (TAC)_12_ substrates (See also Supplementary Fig. 11). The data and error estimate were derived from three independent measurements. **(f)** eRRBS analysis revealing relative methylation of the indicated context in the TKO cells rescued with WT hDNMT3B or N656I (depth of reads >10 and p < 0.0001) **(g**,**h)** Close-up view of the protein-DNA contacts at the RD interfaces of DNMT3A in the previously reported DNMT3A-DNA complex (PDB 5YX2) (**g**) and DNMT3B in the DNMT3B-DNA complex (**h**). **(i)** Preference of mDNMT3B mutants for the SatII substrate analyzed by radioactive methylation assays. Data are displayed as average values in relation to WT DNMT3B. Error bars display standard deviations based on two independent experiments. m DNMT3B residues G828, G830 and K834 correspond to hDNMT3B G822, G824 and K828 (see Supplementary Fig. 10). For recognition of the SATII substrate by WT DNMT3B, refer to Fig. 1f. Statistical analysis used two-tailed Student’s t-test for the difference from WT: ***, *p* < 0.001.

Our NGS analysis revealed an about two-fold higher level of non-CpG methylation for DNMT3B when compared with DNMT3A (Fig. 4d), consistent with previous reports on their differential CpG/CpA specificity^25, 28^. To examine whether the conformational divergence between the catalytic loops of DNMT3A and DNMT3B impact their CpG specificities, we measured the enzymatic activities of DNMT3A and DNMT3B, WT and mutants, on a DNA duplex containing either (GAC)_12_, (AAC)_12_ or (TAC)_12_ repeats. Consistent with the NGS analysis, WT DNMT3B shows 2-4 fold lower CpG/CpA and CpG/CpT specificities than WT DNMT3A (Fig. 4e and Supplementary Fig. 11). Strikingly, swapping this DNMT3B-specific H-bond inverted the CpG specificities of DNMT3A and DNMT3B: the DNMT3B N656I mutation increased the CpG specificity by 2.0 to 1.1-fold, whereas the DNMT3A I715N mutation decreased the CpG specificity by 1.7 to 1.2-fold (Fig. 4e and Supplementary Fig. 11). Consistently, our eRRBS-based methylome data obtained from the WT and N656I-reconstituted TKO cells (two replicates per group; Supplementary Fig. 12a-c and Supplementary Table 4) show that, the DNMT3B N656I mutant displayed a ∼2.6- and 1.4-fold decrease in the relative activity for CpA/CpG and CpT/CpG, respectively, when compared with WT DNMT3B (Fig. 4f). Together, these data suggest that the divergent conformational dynamics of the catalytic loop attributes to the differential CpG/CpH specificities of DNMT3A and DNMT3B.

### The homodimeric RD interfaces of DNMT3A and DNMT3B diverge on DNA binding

Comparison of the DNMT3B-DNA and DNMT3A-DNA complexes also reveals a marked difference in the DNA conformation: In comparison with the DNMT3A-bound DNA, the DNMT3B-bound DNA shows a sharper kink of the central segment arching over the RD interface, bending further away from the protein (Supplementary Fig. 10a). This conformational difference reflects the distinct protein-DNA contacts on a weakly conserved segment of the RD interface (residues G822-K828 of DNMT3B and S881-R887 of DNMT3A; Supplementary Fig. 10b): While in the DNMT3B complex only R823 and G824 form hydrogen bonds with DNA residues, in the DNMT3A complex, S881, R882, L883 and R887 all engage hydrogen bonding or van der Waals contacts with the DNA backbone, explaining the closer approach of the DNA to the protein in the DNMT3A complex (Fig. 4g,h). Along the line, mutation of G822, G824 and K828 in DNMT3B into their corresponding residues in DNMT3A resulted in a markedly reduced preference toward the SatII substrate (Fig. 4i and Supplementary Fig. 13), indicating that the differential flanking sequence preferences of DNMT3B and DNMT3A and the preference of DNMT3B for the SatII target are partially encoded in the DNA-contacting residues in the RD interface.

### Crystal structure of the DNMT3B-CAG complex

To gain insight into the structural basis of DNMT3B-mediated CpA methylation, we also determined the crystal structure of DNMT3B-DNMT3L tetramer covalently bound to a DNA duplex containing two CAG motifs (DNMT3B-CAG complex) (Fig. 5a,b). The structure of the DNMT3B-CAG complex resembles that of the DNMT3B-CpG complexes with subtle conformational differences: the replacement of the CpG dinucleotide with CpA leads to the loss of the base-specific contacts of N779 with the bases next to the methylation site (Fig. 5c), as well as a reduced packing between the A6 base and the catalytic loop (Fig. 5d,e). On the other hand, K777 forms a side-chain hydrogen bond with the N7 atom of G7 (Fig. 5c), providing an explanation for DNMT3B’s preference for a G at the +1 site in the context of non-CpG DNA. In addition, the water-mediated hydrogen bond between DNMT3B T775 and the N7 atom of G6, observed in the DNMT3B-CGA complex (Fig. 3d), and in the corresponding sites of the DNMT3A-DNA complex^33^, is preserved in the DNMT3B-CpA DNA complex now involving the N7 atom of A6 (Fig. 5c), which may contribute to the fact that CpA represents the next most favorable methylation site of DNMT3B and DNMT3A, following CpG.

**Figure 5.**
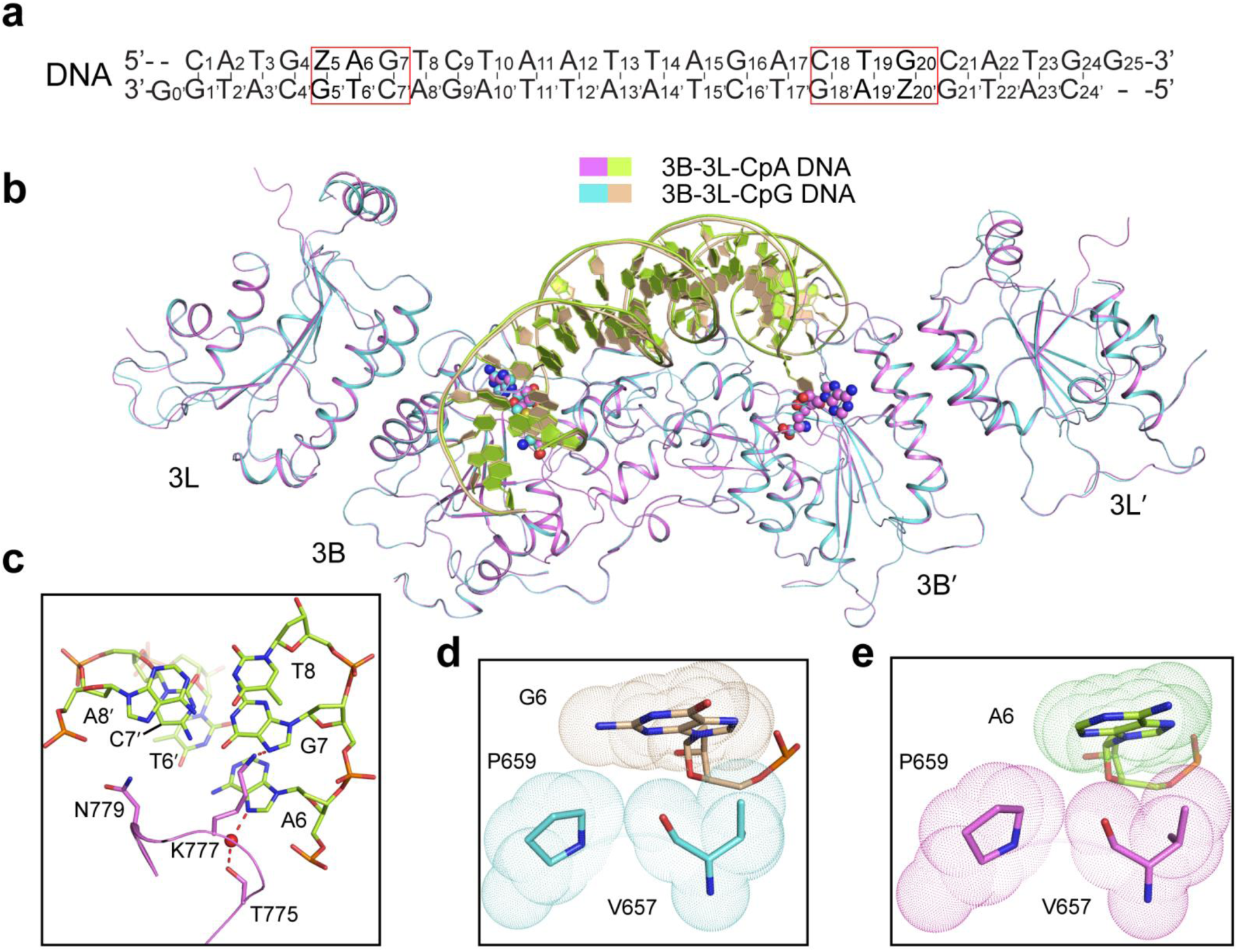
Structure of the DNMT3B-DNMT3L tetramer in complex with CAG DNA. **(a)** DNA sequence used for the structural study. **(b)** Structural overlay of the DNMT3B-CAG DNA and DNMT3B-CGA DNA complexes. **(c)** Close-up view of the interactions between the TRD loop of DNMT3B and the CAG DNA. **(d**,**e)** Close-up view of the van der Waals contacts between the catalytic loop of DNMT3B and G6 in the ZpG site of the CGA complex (d) or A6 in the ZpA site of the CAG complex (e).

### DNMT3B K777 recognizes the +1 flanking nucleotide of the methylation site

Our previous study on the DNMT3A-DNA complex revealed that a hydrogen bond formed between residues R882 and S837 of DNMT3A provides an intramolecular link between the RD interface and the TRD loop upon DNA binding (Fig. 6a)^33^. Interestingly, this interaction is abolished in DNMT3B, because R823 adopts a different conformation and DNA-binding mode than its DNMT3A counterpart R882 (Fig. 6b). Accordingly, the TRD loops of DNMT3B and DNMT3A adopt distinct side-chain conformations for CpG recognition (Fig. 6c): In the DNMT3A-CGT complex, R836 donates a hydrogen bond to the Gua6-O6 atom, whereas the corresponding residue K777 in DNMT3B points toward the +1 flanking nucleotide; instead, in the DNMT3B-CpG complexes the Gua6-O6 atom receives a hydrogen bond from N779, unlike the corresponding N838 in the DNMT3A-CGT complex that does not make any base-specific contact with the CpG site (Fig. 6a,c)^33^. Structural comparison of the three DNMT3B-CpA/CpG complexes reveals distinct major groove environments and side-chain conformations of K777 (Fig. 6d-i and Supplementary Fig. 14a-c): in the DNMT3B-CAG complex, K777 is oriented toward the +1 nucleotide G7 to form a base-specific hydrogen bond, as described above, as well as an electrostatic interaction with the backbone phosphate of G7 (Fig. 6e); likewise, in the DNMT3B-CGA complex, K777 points toward A7 to engage van der Waals contacts and a water-mediated hydrogen bond with the DNA backbone (Fig. 6g); in contrast, in the DNMT3B-CGT complex, in which A7 was replaced by thymine, the side chain of K777 moves away from the DNA backbone, presumably resulting from the reduction in major groove depth by the base ring of T7 (Fig. 6i,j). These observations suggest that DNMT3B K777 reads a combined feature of shape and polarity of the +1 flanking base, distinct from its counterpart in DNMT3A (R836), which recognizes the CpG site in the CGT complex directly through hydrogen-bonding interactions (Fig. 6a)^33^. To further determine the role of DNMT3B K777 in DNA recognition, we generated the K777A-mutated DNMT3B-DNMT3L tetramer in complex with the CGT DNA and determined the structure at 2.95 Å resolution (Supplementary Fig. 14d). The K777A structure aligns well with that of the WT DNMT3B-CGT complex (Supplementary Fig. 14d), but exhibits a subtle conformational change along the G6-T7 step: in comparison with the WT DNMT3B-CGT complex, the T7 base in the K777A DNMT3B-CGT complex undergoes a slide movement toward the major groove, further reducing the groove depth around the +1 site (Fig. 6k,l). Consequently, the G6-T7 step appears to engage a stronger base-stacking interaction in the K777A DNMT3B-CGT complex (Fig. 6m) than in the WT DNMT3B-CGT complex (Fig. 6n). It is worth mentioning that the hydrogen bond between DNMT3B N779 and the CpG site is unaltered by the K777A mutation (Supplementary Fig. 14e). Likewise, structural comparison of the previously reported WT and R836A-mutated DNMT3A-CGT complexes^33^ reveals that introduction of the R836A mutation does not lead to a conformational change of neighboring N838 (Supplementary Fig. 14f). These data suggest that the conformational divergence between the TRD loops of DNMT3A and DNMT3B in the CGT complexes is unlikely attributed to the R-to-K change at the DNMT3B R836 and DNMT3A K777 sites. Together, these structural studies reveal a distinct interplay of the RD interface and TRD loop between DNMT3A and DNMT3B, suggesting a role of DNMT3B K777 in shaping the preference of DNMT3B toward the flanking sequence of the CpG site.

**Figure 6.**
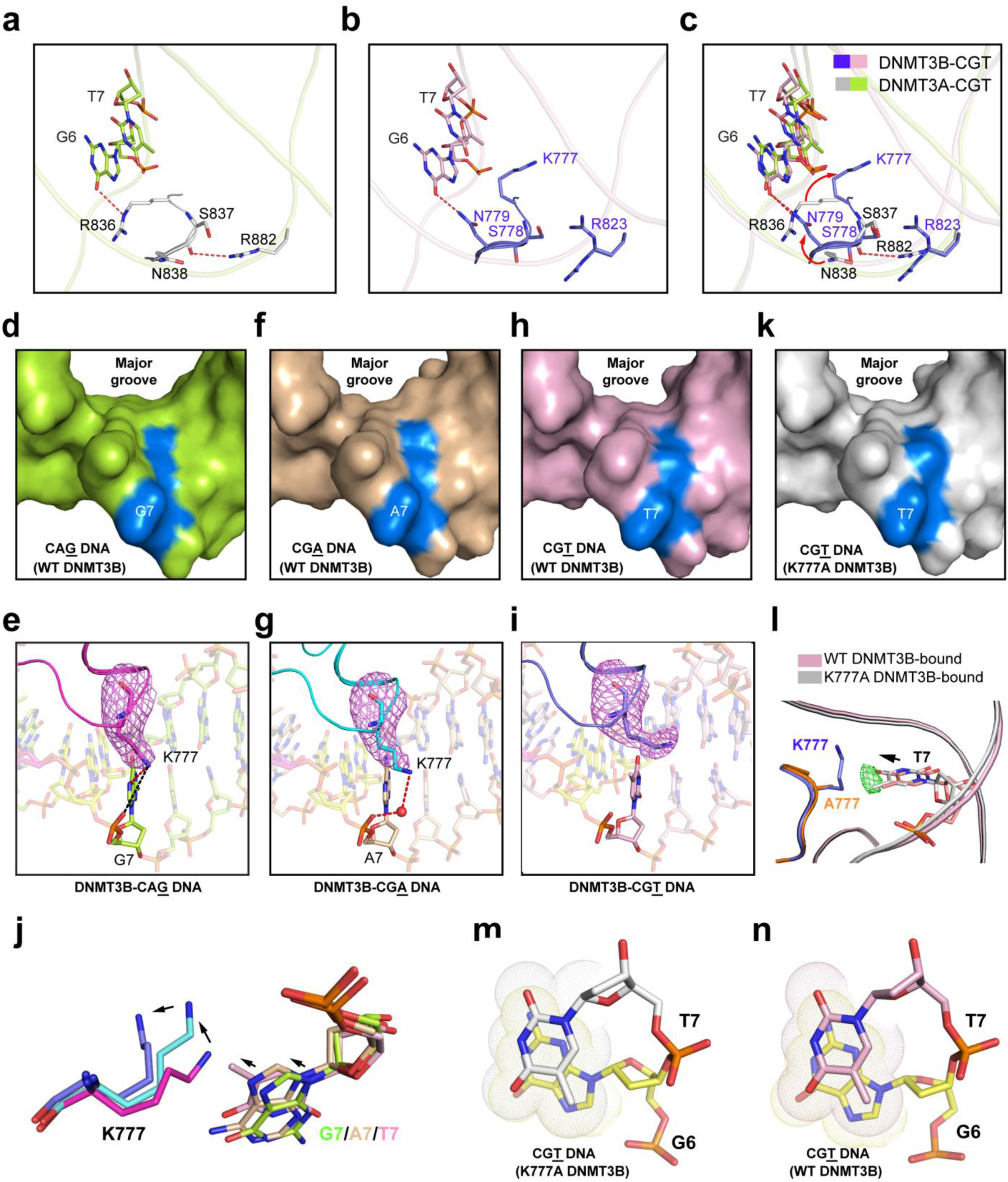
Recognition of the DNA shape of the +1 flanking site by DNMT3B K777. **(a-c)** Close-up comparison of the protein-DNA contacts at the TRD loops of hDNMT3A (b) and hDNMT3B (c) in their respective DNA complexes. **(d-i)** Surface and stick representations of the CAG (**d**,**e**), CGA (**f**,**g**) and CGT (**h**,**i**) DNAs bound to the DNMT3B-DNMT3L tetramer. The +1 nucleotides are colored in marine in (**d**), (**f**) and (**h**). Fo-Fc omit map of the K777 (light magenta) is contoured at 2.0 σ level in (**e**), (**g**) and (**i**). The hydrogen bonding and electrostatic interactions are shown as red and black dashed lines, respectively. The water molecule is shown as red sphere. (**j**) Structural superposition of DNMT3B K777 and interacting +1 nucleotides in CAG, CGA and CGT DNA complexes. The color schemes are the same as in (**e**), (**g**) and (**i**). The relative positioning of the +1 bases and DNMT3B K777 are indicated by arrows. (**k**) Surface view of the CGT DNA bound to K777A-mutated DNMT3B-DNMT3L tetramer, with the +1 nucleotide T7 colored in marine. (**l**) Structural comparison of the +1 nucleotide (T7) at the CpG site of the DNA bound to WT or K777A-mutated DNMT3B. The T7 nucleotides associated with WT and K777A-mutated DNMT3B-CGT complexes are colored in light pink and silver, respectively. The conformational shift of the T7 nucleotide between the two complexes is indicated by arrow as well as the Fo-Fo difference map of T7 (green; contoured at 2.0 σ level) derived from the electron density for the WT and K777A DNMT3B complexes. (**m**,**n**) Close-up view of the stacking interaction between CpG guanine (G6) and +1 base (T7) of CGT DNA bound to K777A-mutated (**m**) or WT (**n**) DNMT3B-DNMT3L.

### DNMT3B K777 mediates the +1 flank site preference of DNMT3B

To analyze the effect of K777 on the substrate preference of DNMT3B, we measured the enzymatic activities of DNMT3B-DNMT3L, WT and mutants, on hemimethylated DNAs containing unmethylated (GTC)_12_ or (GAC)_12_ on one strand while methylated (GA^m^C)_12_ or (GT^m^C)_12_ on the complimentary strand, referred to as CGT and CGA DNAs, respectively. WT DNMT3B shows a ∼2-fold preference for CGA DNA over CGT DNA (Fig. 7a); in contrast, the K777A mutant shows increased methylation for both CGT and CGA DNA, but with preference for CGT over CGA (Fig. 7a), which likely arises from the altered base stacking interaction between the +1 base and the CpG guanine (Fig. 6m,n). The N779A mutation led to reduced activity toward CGA (Fig. 7a), but not CGT, consistent with its role in contacting the +1 T on the non-target strand in the CGA complex (Fig. 3d). Likewise, our NGS analysis reveals that the pronounced preference of DNMT3B for non-CpG methylation at sites flanked by G at the +1 site was completely lost in K777A, which instead showed a preference for T at this site in both CpG and non-CpG context (Fig. 7b). Finally, we have compared the cellular methylation profiles obtained from TKO cells reconstituted with WT versus K777A-mutated DNMT3B (two replicates per group, see Supplementary Table 5 and Supplementary Fig. 15a-e). Similar to the *in vitro* analysis, the K777A mutation led to a G-to-T preference change at the +1 flanking site in all sequence contexts, relative to WT DNMT3B (Fig. 7c). In contrast, eRRBS analysis of TKO cells reconstituted with the N779A mutant (Supplementary Table 5 and Supplementary Fig. 15a-e) showed that this mutant retained a preference for a G at the +1 flanking site in the context of non-CpG DNA, resembling what was observed for WT DNMT3B (Fig. 7d). Nevertheless, the N779A mutation led to a modest decrease in the CpG specificity *in vitro* and in cells (Supplementary Fig. 16a-c), in line with its role in CpG recognition. These findings establish that K777 of DNMT3B functions as a crucial determinant sensing different sequence contexts flanking the methylation site, which is distinctive from what was observed for DNMT3A.

**Figure 7.**
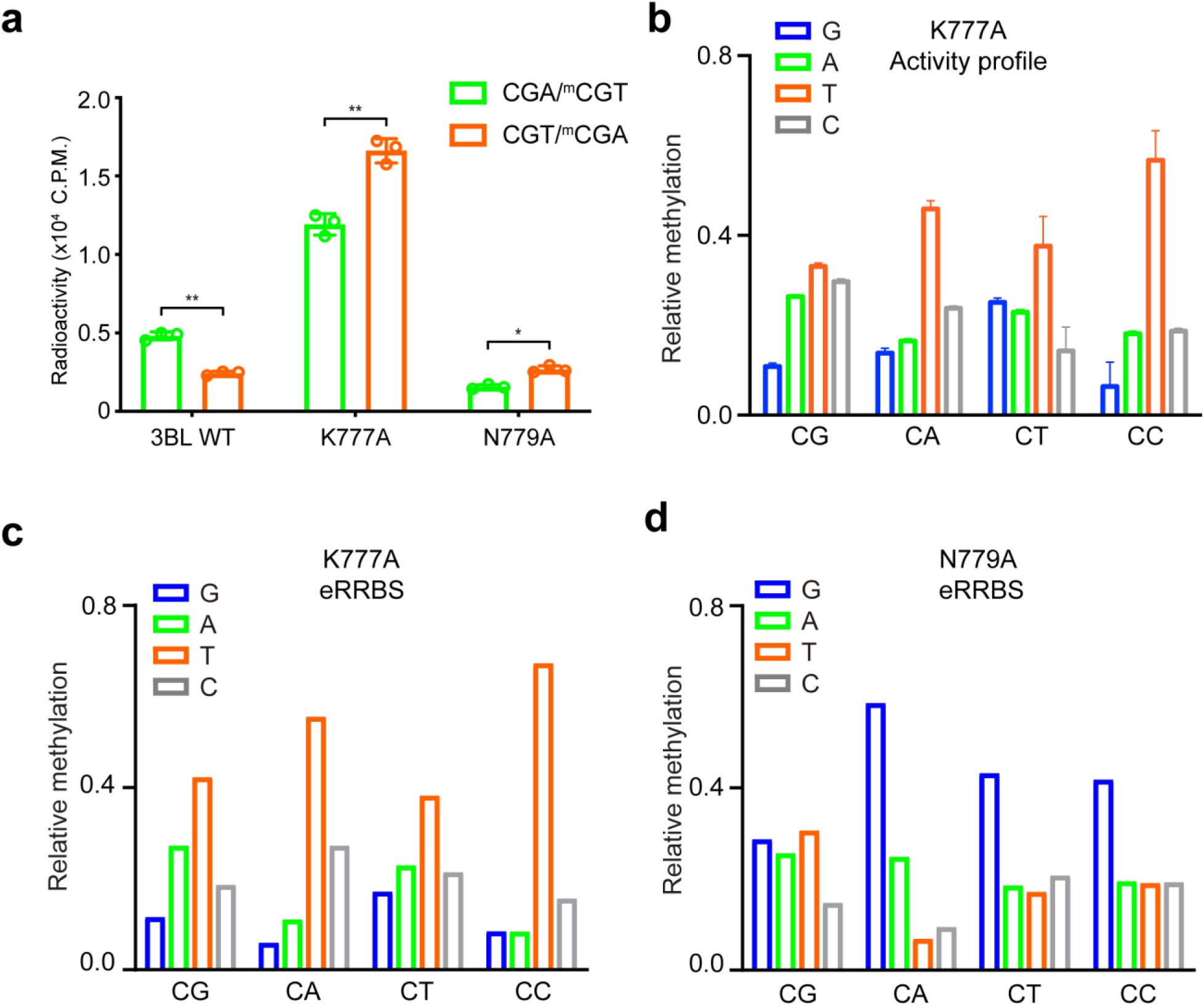
K777 mediates the +1 flanking site preference of DNMT3B. **(a)** *In vitro* methylation analysis of hDNMT3B-hDNMT3L, WT or mutants, on hemimethylated CGT and CGA DNA. The data and error estimate were derived from three independent measurements. Statistical analysis used two-tailed Student’s t-test. *, *p* < 0.05, **, *p* < 0.01. **(b)** Deep enzymology profiles for CpG and non-CpG methylation of hDNMT3B K777A averaged for the different +1 flanking base pairs. The data show average methylation levels based on two independent experiments. Error bars indicate the standard deviation. **(c**,**d)** eRRBS-based methylation analysis of CpG and non-CpG methylation of DNMT3B K777A (**c**) and DNMT3B N779A (**d**) averaged for the different +1 flanking base pairs (depth of reads >10 and *p* <0.0001).

## Discussion

The functional interplay between DNMT3A and DNMT3B is critical for the dynamic programming of epigenetic regulation in development and disease. Through comprehensive structural, biochemical and enzymatic characterizations of DNMT3B and DNMT3A in CpG and non-CpG methylation, this study provides an unprecedented view on DNMT3B- and DNMT3A-mediated *de novo* methylation, explaining their distinct functionalities in cells.

The deep enzymology approach developed in this study, combined with the eRRBS-based cellular methylome profiling, allowed high-resolution characterization of the flanking sequence preferences of DNMT3A and DNMT3B, which manifest >100-fold different methylation rates of CpG sites across different sequence contexts. Importantly, our data demonstrate that the enzymatic divergence between DNMT3A and DNMT3B contributes to their differential activities on SatII repeats, providing one molecular explanation for why hypomethylation of SatII sequences is strongly connected with DNMT3B mutations, and DNMT3A cannot compensate for a lack of DNMT3B activity. The flanking sequence preferences of DNMT3A and DNMT3B also explain the previously observed preferences of both enzymes in mouse ES cells, where DNMT3B methylates minor satellite repeats and DNMT3A methylates major satellite repeats^17^.

Importantly, this study reveals a multi-layered substrate recognition mechanism. First, the formation of a specific H-bond in the catalytic loop of DNMT3B gives rises to a reduced CpG/CpH specificity in DNMT3B, likely due to a conformational stabilization effect. Second, the different protein-DNA interactions at the RD interface of DNMT3B and DNMT3A diversify their intramolecular interaction between the RD interface and TRD, which triggers changes in the side-chain conformations of TRD loop residues, including DNMT3B K777, S778 and N779, leading to a distinct TRD-DNA interaction of DNMT3B. Third, this study reveals a role for the TRD loop in fine-tuning substrate readout of DNMT3B. Strikingly, K777 undergoes significant side-chain conformational changes in response to changes in the major groove environments caused by different +1 bases, providing a mechanism for readout of +1 nucleotides by DNMT3B. This base shape-directed readout of the +1 flanking site by DNMT3B K777 on one hand may reduce the requirement for the N779-CpG contact when the CpG substrate is in a “favorable” flanking sequence context, on the other hand, shifts the preference of DNMT3B toward G at the +1 site on non-CpG substrates. This study therefore adds a new example to the growing family of DNA-interacting proteins that recognizes the DNA shape as a readout mechanism^44^.

This study provides the molecular basis underlying the sequence preferences of DNMT3A and DNMT3B on the +1 flanking site. How other flanking sites (e.g. the −2 site, in which a preference toward T was observed for both DNMT3A and DNMT3B) affect DNMT3A- and DNMT3B-mediated methylation remains to be determined. It is also worth mentioning that the genomic targeting of DNMT3A and DNMT3B in cells are further regulated by additional factors, including their N-terminal domains and DNMT3L. For instance, a previous study has demonstrated that DNMT3L modulates cellular *de novo* methylation activities through focusing the DNA methylation machineries on well-chromatinized DNA templates^32^. In addition, despite that structural analysis of both DNMT3B-DNA and DNMT3A-DNA complexes reveals that DNMT3L does not directly engage in the DNA interaction, the role of DNMT3L in stimulating the enzymatic activity and/or stability of DNMT3A and DNMT3B has been well established^6-8, 42, 45^, which may attenuate the impact of the intrinsic sequence specificities of these enzymes on the landscape of DNA methylation^32^. How the intrinsic preferences of DNMT3A and DNMT3B interplay with other cellular factors in regulating genomic DNA methylation awaits further investigation.

## Acknowledgments

We would like to thank staff members at the Advanced Light Source (ALS), Lawrence Berkeley National Laboratory for access to X-ray beamlines. This work was supported by NIH grants (1R35GM119721 to J.S.; 1R01CA215284 and 1R01CA211336 to G.G.W., and 5R21ES025392 to Y.W.), the Intramural Research Program of the National Institute of Environmental Health Sciences, NIH (ES101965 to P.A.W.), and the Deutsche Forschungsgemeinschaft DFG (grants JE 252/10 and JE 252/36 to A.J.). We are also grateful for professional assistance of UNC facilities including Genomics Core, which are in part supported by the UNC Cancer Center Core Support Grant P30-CA016086. G.G.W. is an American Cancer Society (ACS) Research Scholar and a Leukemia & Lymphoma Society (LLS) Scholar.

## Author Contributions

LG, ZMZ and JL performed the structural studies. LG, WR and HA performed *in vitro* enzymatic assays with DNMT3A-DNMT3L and DNMT3B-DNMT3L proteins. JS directed the structural and enzymatic study for DNMT3A-DNMT3L and DNMT3B-DNMT3L and analyzed the data. ME conducted enzyme purification and enzyme kinetics and deep enzymology methylation reactions with DNMT3B and DNMT3A with support from MD and RJ. MD performed and analyzed the deep enzymology experiments with K777A. SA, PB and AJ developed and implemented the deep enzymology workflow with DNMT3A and DNMT3B, conducted the experiments and analyzed the data. DC and YG carried out eRRBS and DNMT3A- and DNMT3B-mediated genomic methylation studies under guidance of GGW. JY performed LC-MS measurement of global cytosine methylation under the guidance of YW. HU and SAG analyzed eRRBS methylome data under direction of PAW and GGW. GGW, AJ and JS prepared the manuscript with input from all authors.

## Author Information

The authors declare no competing financial interests. Correspondence and requests for materials should be addressed to G.G.W., A.J. or J.S.

## Data availability

Coordinates and structure factors for the DNMT3B complexes have been deposited in the Protein Data Bank under accession codes 6U8P, 6U8V, 6U8X and 6U8W. The eRRBS data have been deposited in Gene Expression Omnibus (GEO).

## Code availability

The custom scripts used for data analysis are available upon request.

## Methods

### Protein expression and purification

The MTase domains of human DNMT3B (residues 562–853 of NCBI accession NM_006892) or DNMT3A (residues 628–912 of NCBI accession NM_022552) were co-expressed with the C-terminal domain of human DNMT3L (residues 178–386 of NCBI accession NM_175867) on a modified pRSFDuet-1 vector (Novagen), in which the DNMT3B or DNMT3A sequence was preceded by a hexahistidine (His_6_) and SUMO tag. The *Escherichia coli* BL21 DE3 (RIL) cell strains transformed with the DNMT3A- or DNMT3B-expression plasmids were first grown at 37 °C, but shifted to 16 °C after induction by IPTG at an OD_600_ (optical density at 600 nm) of 0.8. The cells continued to grow overnight. The His_6_-SUMO–tagged DNMT3B or DNMT3A fusion proteins in complex with DNMT3L were purified using a Ni^2+^-NTA column. Subsequently, the His_6_-SUMO tag was removed through Ubiquitin-like-specific protease 1 (ULP1) cleavage, followed by ion exchange chromatography using a Heparin HP column (GE Healthcare) and further purification through size-exclusion chromatography on a HiLoad 16/600 Superdex 200 pg column (GE Healthcare) in a buffer containing 20 mM Tris-HCl (pH 8.0), 100 mM NaCl, 0.1% β-mercaptoethanol, and 5% glycerol. Purified protein samples were stored in −80°C for future use. For DNMT3-alone methylation experiments the catalytic domains of murine DNMT3A (residues 608-908 of NM_001271753) and DNMT3B (residues 558-859 of NM_001271744) were expressed and purified as described ^26, 47^.

To generate the covalent protein-DNA complex, a 25-mer Zebularine-containing DNA (CGA: 5′-CAT GZG ATC TAA TTA GAT CGC ATGG-3′, CGT: 5′-GCA TGZ GTT CTA ATT AGA ACG CATG-3′, CAG: 5′-CAT GZA GTC TAA TTA GAC TGC ATGG-3′, Z: zebularine) was self-annealed and incubated with WT or mutant DNMT3B-DNMT3L in 20 mM Tris-HCl (pH 8.0), 20% glycerol, and 40 mM DTT at room temperature. The reaction products were further purified through a HiTrap Q XL column (GE Healthcare), followed by size-exclusion chromatography on a HiLoad 16/600 Superdex 200 pg column. The purified covalent protein-DNA complexes were concentrated to about 0.1– 0.2 mM using Ultracel®-10K Centrifugal Filters (Millipore) in a buffer containing 20 mM Tris-HCl (pH 8.0), 100 mM NaCl, 0.1% β-mercaptoethanol and 5% glycerol.

### Crystallization and structure determination

The crystals of covalent complexes of WT or mutant DNMT3B-DNMT3L-DNA complexes were generated by the hanging-drop vapor-diffusion method at 4°C from drops mixed from 0.5 μL of the protein solution and 0.5 μL of precipitation solution containing 0.1 M Tris-HCl (pH 8.0), 200 mM MgCl_2_, 8% PEG4000 and 0.2 mM AdoHcy. Crystals for the DNMT3B-DNMT3L tetramer in complex with the CGT were generated by the hanging-drop vapor-diffusion method at 16 °C, from drops mixed from 0.5 μL of DNMT3B-DNMT3L-DNA solution and 0.5 μL of precipitant solution containing 0.1 M Tris-HCl (pH 8.0), 100 mM MgCl_2_, 7% PEG8000 and 0.2 mM AdoHcy. All crystals were soaked in cryo-protectant made of the precipitation solution supplemented with 25% glycerol, before flash frozen in liquid nitrogen for X-ray data collection. The X-ray diffraction datasets for the DNMT3B-DNMT3L-DNA complexes were collected at selenium peak wavelength on the BL 5.0.1 or BL 5.0.2 beamlines at the Advanced Light Source, Lawrence Berkeley National Laboratory. The diffraction data were indexed, integrated, and scaled using the HKL 2000 program^48^. The structures of the complexes were solved by molecular replacement method using PHASER^49^, with the structure of DNMT3A-DNMT3L-DNA complex (PDB 5YX2) as a search model. Further modelling of the covalent DNMT3B-DNMT3L-DNA complexes was performed using COOT^50^ and subjected to refinement using the PHENIX software package^51^. The same R-free test set was used throughout the refinement. The statistics for data collection and structural refinement of the productive covalent DNMT3B-DNMT3L-DNA complexes are summarized in Table 1.

### *In vitro* DNA methylation assays

DNA methylation kinetics shown in Fig. 3f, 4e, 7a and Supplementary Fig. 16c were conducted using the C-terminal domains of DNMT3A-DNMT3L or DNMT3B-DNMT3L. A 20 μL reaction mixture contained 0.75 µM DNA, 0.3 µM DNMT3B-DNMT3L or DNMT3A-DNMT3L tetramer, 2.5 μM *S*-adenosyl-L-[methyl-^3^H]methionine (specific activity 18 Ci/mmol, PerkinElmer) in 50 mM Tris-HCl, pH 8.0, 0.05% β-mercaptoethanol, 5% glycerol and 200 μg/mL BSA. The DNA methylation assays were carried out in triplicate at 37°C for 40 min before being quenched by addition of 5 µL of 10 mM unlabeled SAM. For detection, 12.5 µL of reaction mixture was spot on DEAE Filtermat paper (PerkinElmer) and dried. The DEAE paper was then washed sequentially with 3 x 5 mL of cold 0.2 M ammonium bicarbonate, 5 mL of Milli Q water, and 5 mL of ethanol. The DEAE paper was air dried and transferred to scintillation vials filled with 5 mL of ScintiVerse (Fisher). The radioactivity of tritium was measured with a Beckman LS6500 counter.

For experimental validation of the SatII preference of DNMT3B, the 6-bp SatII flanking sequence (TCCATTCGATGATG) was integrated into a regular 30-mer CpG substrate. As reference the AGGCGCCC substrate which is highly disfavored by DNMT3B was used and embedded in the same overall sequence context. Both substrates were used with a hemimethylated CpG site with methylation in lower strand and with a fully methylated CpG site. In each case, the activity observed with the fully methylated substrate was subtracted from the activity detected with the hemimethylated substrate to specifically determine the methylation of the target CpG in the upper DNA strand^30, 52^.

DNA methylation kinetics shown in Fig. 1f, 4i, and Supplementary Fig. 3, 7a,b, 13 were measured using 1 µM biotinylated double stranded 30-mer oligonucleotides containing a single CpG site basically as described^52^. Using a standard substrate GAG AAG CTG GGA CTT C<u>CG</u> GGA GGA GAG TGC^34^, flanking sequences selected to be preferred or disfavored by DNMT3A and DNMT3B were inserted as described in the text and figure legends. In all substrates, the lower DNA strand was biotinylated. To study the specific methylation of one CpG site in one DNA strand, each different oligonucleotide substrate was used in hemimethylated form and with the central CpG site in fully methylated form and the methylation rate observed with the fully methylated substrate was subtracted from the rate observed with corresponding the hemimethylated one. DNA methylation was measured by the incorporation of tritiated methyl groups from radioactively labeled SAM (Perkin Elmer) into the biotinylated substrate using an avidin–biotin methylation plate assay^47^. The methylation reactions were carried out in methylation buffer (20 mM HEPES pH 7.5, 1 mM EDTA, 50 mM KCl, 0.05 mg/ml bovine serum albumin) at 37°C using 2 µM WT or mutant DNMT3A or DNMT3B catalytic domain. The reactions were started by adding 0.76 µM radioactively labelled SAM. The initial slope of the enzymatic reaction was determined by linear regression. In Fig. 4i, as reference substrate a biotinylated 509-mer DNA containing 58 CpG sites was used at a concentration of 100 nM^47, 52^.

### Deep enzymology experiments

Single-stranded DNA oligonucleotides used for generation of double stranded substrates with CpH or methylated CpG sites embedded in a 10 nucleotide random context were obtained from IDT. The second strand synthesis was conducted by a primer extension reaction using one universal primer. The obtained mix of double-stranded DNA oligonucleotides was methylated by murine DNMT3A or DNMT3B catalytic domain for 60 min at 37 °C in the presence of 0.8 mM S-adenosyl-L-methionine (Sigma) in reaction buffer (20 mM HEPES pH 7.5, 1 mM EDTA, 50 mM KCl, 0.05 mg/mL bovine serum albumin). Reactions were stopped by shock freezing in liquid nitrogen, then treated with proteinase K for 2 hours. Afterwards, the DNA was digested with the BsaI-HFv2 enzyme and a hairpin was ligated using T4 DNA ligase (NEB). The DNA was bisulfite converted using EZ DNA Methylation-Lightning kit (ZYMO RESEARCH) according to the manufacturer protocol, purified and eluted with 10 µL ddH2O.

Libraries for Illumina Next Generation Sequencing (NGS) were produced with the two-step PCR approach. In the first PCR, 2 µL of bisulfite-converted DNA were amplified with the HotStartTaq DNA Polymerase (QIAGEN) and primers containing internal barcodes using following conditions: 15 min at 95 °C, 10 cycles of 30 sec at 94 °C, 30 sec at 50 °C, 1 min and 30 sec at 72 °C, and final 5 min at 72 °C; using a mixture containing 1x PCR Buffer, 1x Q-Solution, 0.2 mM dNTPs, 0.05 U/µL HotStartTaq DNA Polymerase, 0.4 µM forward and 0.4 µM reverse primers in a total volume of 20 µL. In the second PCR, 1 µL of obtained products were amplified by Phusion Polymerase (Thermo) with another set of primers to introduce adapters and indices needed for NGS (30 sec at 98 °C, 10 cycles - 10 sec at 98 °C, 40 sec at 72 °C, and 5 min at 72 °C). PCRII was carried out in 1x Phusion HF Buffer, 0.2 mM dNTPs, 0.02 U/µL Phusion HF DNA Polymerase, 0.4 µM forward and 0.4 µM reverse primers in a total volume of 20 µL. Obtained libraries were pooled in equimolar amounts and purified using NucleoSpin^®^ Gel and PCR Clean-up kit (Macherey-Nagel), followed by a second purification step of gel extraction and size exclusion with AMPure XP magnetic beads (Beckman Coulter). Sequencing was performed at the Max Planck Genome Centre Cologne.

Bioinformatics analysis of obtained NGS data was conducted with the tools available on the Usegalaxy.eu server^53^ and with home written programs. Briefly, fastq files were analyzed by FastQC, 3’ ends of the reads with a quality lower than 20 were trimmed and reads containing both full-length sense and antisense strands were selected. Next, using the information of both strands of the bisulfite-converted substrate the original DNA sequence and methylation state of both CpG sites was reconstituted. CpH and CpN data were split into CpG, CpA, CpT and CpC, as appropriate. In each case average methylation levels of each NNCGNN and NNNCGNNN site were determined. Pearson correlation factors were calculated using Excel. Each experiment was performed in two independent repeats. For downstream analysis, DNMT3A data of repeat 1 and the combined data of DNMT3B were used for further analysis, based on their comparable methylation levels. Sequence logos were calculated using Weblogo 3 (http://weblogo.threeplusone.com/).

### Plasmids

The plasmid that contains the isoform 1 of human DNMT3B (DNMT3B1) was purchased from Addgene (cat # 35522). The DNMT3B1 cDNA was then fused to an N-terminal Flag tag by PCR, followed by subcloning into the pPyCAGIZ vector^33^ (a gift of J. Wang). The pPyCAGIZ construct for expression of the isoform 1 of human DNMT3A (DNMT3A1) was previously described^33^. Point mutation was generated by a QuikChange II XL Site-Directed Mutagenesis Kit (Agilent). All plasmid sequences were verified by sequencing before use.

### Cell lines and tissue culture

The mouse embryonic stem cell line lacking DNMTs (Dnmt1^-/-^ Dnmt3a^-/-^ Dnmt3b^-/-^ or TKO-ESCs; a gift from Dr. M. Okano) were cultivated as previously described^33^. 3KO-ESCs were transfected by Lipofectamine 2000 (Invitrogen) with the pPyCAGIZ empty vector or that carrying WT DNMT3A1, WT DNMT3B1 or its mutant. Transduced ES cells were selected in culture medium with 50μg/mL Zeocin (Invitrogen) for over two weeks, following by establishment of pooled stable-expression cell lines and independent single-cell-derived clonal lines as described before^33^.

### Antibodies and western blotting

Antibodies used for immunoblotting include α-Flag (Sigma; M2), DNMT3B (Santa Cruz bio., G-9: sc-376043) and α-Tubulin (Sigma). Immunoblotting was conducted as previously described^33^.

### Quantification of 5-methyl-2′-deoxycytidine (5mC) in genomic DNA

Enzymatic digestion of cellular DNA and LC-MS/MS measurements of the levels of 5-mdC in the resulting nucleoside mixture was carried out as described previously^33^.

### eRRBS and data analysis

eRRBS analysis of different cell samples was carried out in parallel to generate datasets for across-sample comparisons (such as WT DNMT3A1 versus DNMT3B1, or WT vs. mutant). Specifically, eRRBS was performed after DNA cleavage with restriction enzymes of MspI, BfaI, and MseI as described before^33^, followed by deep sequencing with an Illuminia sequencer (High Throughput Genomic Sequencing Facility [HTSF], UNC at Chapel Hill). For data analysis, general quality control checks were performed with FastQC v0.11.2 (http://www.bioinformatics.babraham.ac.uk/projects/fastqc/). The last 5 bases were clipped from the 3’ end of every read due to questionable base quality in this region, followed by filtration of the sequences to retain only those with average base quality scores of more than 20. Examination of the 5’ ends of the sequenced reads indicated that 70%-90% (average 83.0%) were consistent with exact matches to the expected restriction enzyme sites (i.e. MspI, BfaI, and MseI). Approximately 85% of both ends were consistent with the expected enzyme sites. Adapter sequence was trimmed from the 3’ end of reads via Cutadapt v1.2.1 (parameters -a AGATCGGAAGAG -O 5 -q 0 -f fastq; https://doi.org/10.14806/ej.17.1.200). Reads shorter than 30 nt after adapter-trimming were discarded. Filtered and trimmed datasets were aligned via Bismark v0.18.1 (parameters -X 1000 --non_bs_mm) ^54^ using Bowtie v1.2^55^ as the underlying alignment tool. The reference genome index contained the genome sequence of enterobacteria phage λ (NC_001416.1) in addition to the mm10 reference assembly (GRCm38). For all mapped read pairs, the first 4 bases at the 5’ end of read1 and the first 2 bases at the 5’ end of read2 were clipped due to positional methylation bias, as determined from QC plots generated with the ‘bismark_methylation_extractor’ tool (Bismark v0.18.1). To avoid bias in quantification of methylation status, any redundant mapped bases due to overlapping read ends from the same read pair were trimmed. Read pairs in which either read end had 3 more or methylated cytosines in non-CpG context were assumed to have escaped bisulfite conversion and were discarded. Finally, mapped read pairs were separated by genome (mm10 or phage λ). Read pairs mapped to phage λ were used as a QC assessment to confirm that the observed bisulfite conversion rate was >99%. Read pairs mapped to the mm10 reference genome were used for downstream analysis. Although the eRRBS data does carry stranded information, data from the plus and minus strands have been collapsed in this analysis.

Calling of methylated cytosine was performed with Bismark. The non-conversion rates of produced eRRBS datasets were all less than 0.3% and the statistical significance of methylation at each C site was assessed by binomial testing as described^33^. Following initial analysis of each individual dataset, results from the replicated eRRBS experiments that show good correlation were then combined for following analysis, with only those sites with a coverage of 10 reads or more used to determine the averaged methylation plots shown in main figures.

## Supplementary Information

**Supplementary Figure 1.**
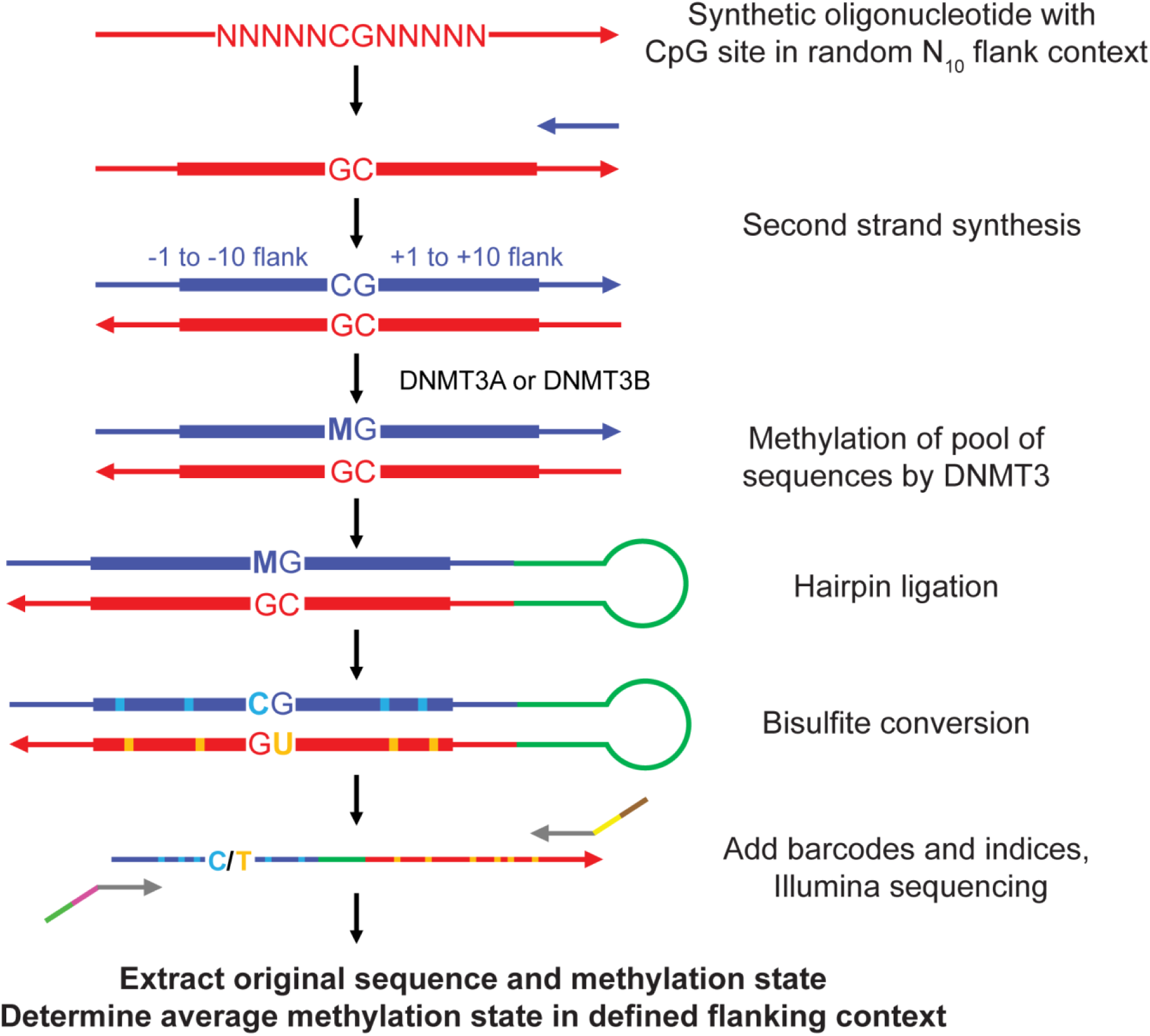
Deep enzymology workflow allowing the methylation of pools of DNA substrates containing target sites in randomized sequence contexts. Pools of DNA substrates with a target CpG site in a randomized flanking context were generated using a synthetic oligonucleotide, which contains one target CpG site flanked by 10 random nucleotides on each side as starting point. In addition, randomized oligonucleotides were used containing the target cytosine followed by 5-methylcytosine (allowing to study the methylation preferences in the context of hemimethylated CpG sites) and in CpH (H=adenine, thymine, or cytosine) or CpN (N=adenine, guanine, thymine, or cytosine) context. Libraries of double stranded DNA substrate molecules containing CpG (or variants thereof) sites in randomized flanking context were then prepared by primer extension. The purified double stranded substrate library was incubated with DNMT3A or DNMT3B in methylation buffer containing AdoMet leading to the methylation of the target sites, depending on the enzymes’ preferences for the respective flanking sequence. After stopping the methylation reaction, a hairpin was ligated to the DNA substrates. Next, bisulfite conversion was carried out which converts cytosine to uracil, but leaves 5-methylcytosine intact. This was followed by amplification with primers specific for the converted DNA. Libraries for Illumina Next Generation Sequencing (NGS) were generated in a two-step PCR approach adding barcodes and indices. After determination of concentrations and validation of the products on acrylamide gels, the products from different methylation reactions were pooled in appropriate ratios and analyzed by Illumina NGS. Data were generated in independent repeats and sequenced at great depth (Supplementary Table 1). Control experiments without enzyme were conducted to determine the efficiency of the bisulfite conversion (Supplementary Table 1). Afterwards the reads were filtered for duplicates and sequencing errors. For each read, the original flank sequence and the methylation state was extracted and compiled in a database. Later average methylation levels in different flanking sequence contexts were extracted, for example in CN, CNN, NNCGNN or NNNCGNNN contexts.

**Supplementary Figure 2.**
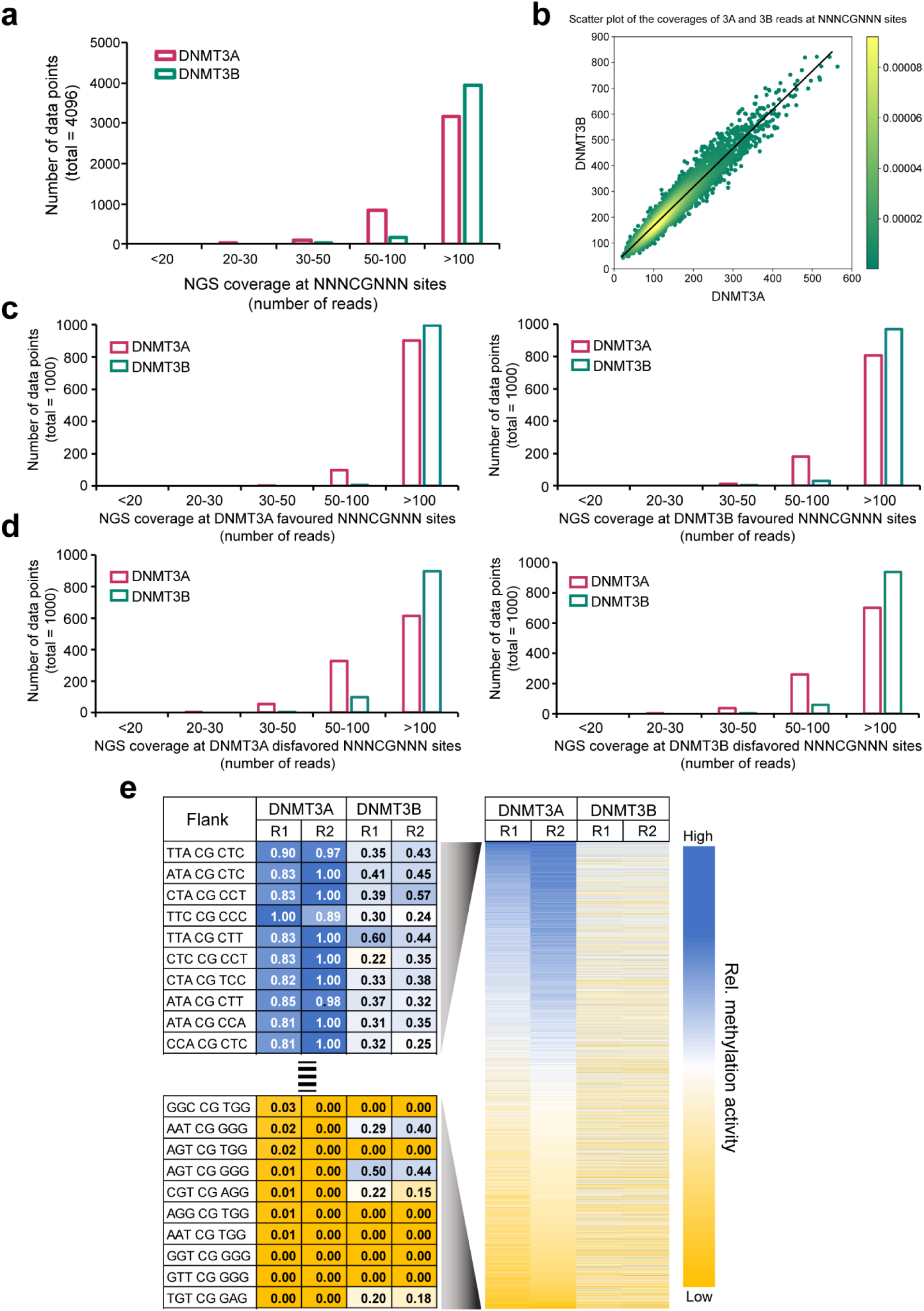
Additional data regarding deep enzymology flanking sequence preferences. **(a**,**b)** Bar (a) and Scatter (b) plots showing the coverage of the NNNCGNNN sites for NGS analysis. **(c)** Bar plots showing the coverage of the top 1000 NNNCGNNN sites most favored by mDNMT3A (left) or mDNMT3B (right). **(d)** Bar plots showing the coverage of the top 1000 NNNCGNNN sites most disfavored by mDNMT3A (left) or mDNMT3B (right). **(e)** Heatmap of the activities of mDNMT3A and mDNMT3B in different NNNCGNNN flanking contexts in two experiments showing the close correlation of the experimental repeats of both data sets and the pronounced differences between mDNMT3A and mDNMT3B. Data are sorted for DNMT3A activity. The left part shows an enlargement of the first and last 20 lines of the heatmap also including the corresponding sequences and normalized activity levels.

**Supplementary Figure 3.**
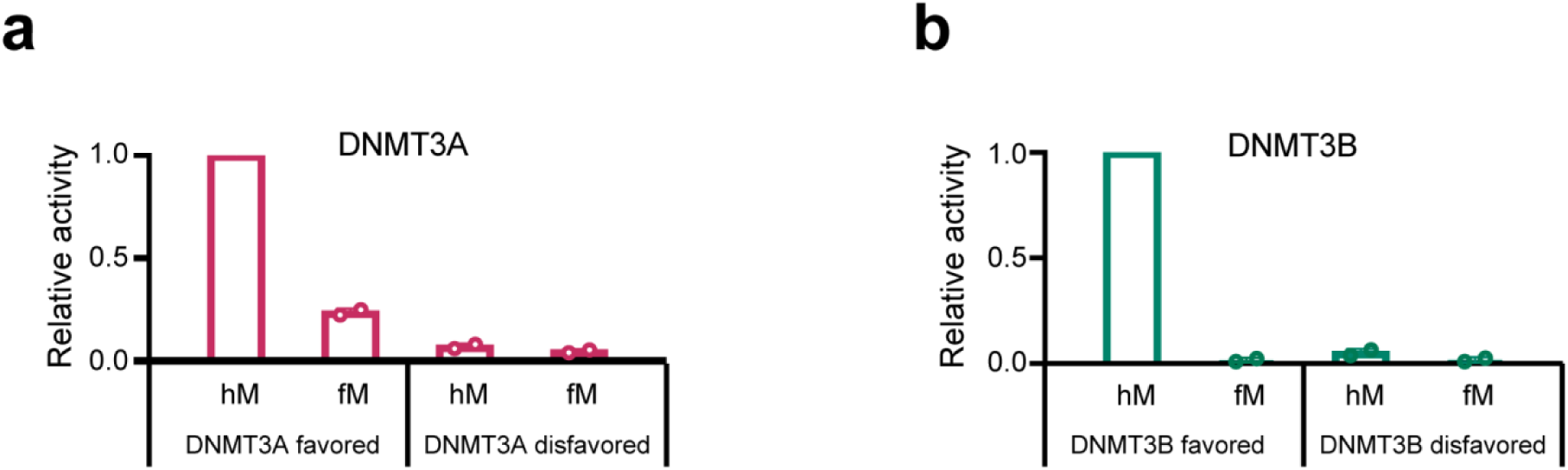
Validation of the deep enzymology experiments. **(a-b)** Enzymatic validation of DNMT3A (a) or DNMT3B (b) on favored and disfavored substrates, using the MTase domains of mDNMT3A and mDNMT3B analyzed by radioactive methylation assays. For both enzymes, the 4096 NNNCGNNN flanks were sorted by preferences (confer Supplementary Fig. 2e). For mDNMT3A, the TTA<u>CG</u>CCC (rank 49) and AGT<u>CG</u>TGG (rank 4014) sites were selected as favored and disfavored substrates, respectively, and subjected to methylation assays. For mDNMT3B, the CTA<u>CG</u>GCT (rank 330) and AGT<u>CG</u>CAA (rank 3328) sites were selected as favored and disfavored substrates. Each substrate was used in hemimethylated form and with fully methylated CpG site to control for methylation of cytosine residues at other places of the substrate. The specific methylation of the CpG in the upper DNA strand is given by the difference of the activities observed on the hemi- and fully methylated substrates. Average values and standard deviations based on two experimental repetitions are displayed.

**Supplementary Figure 4.**
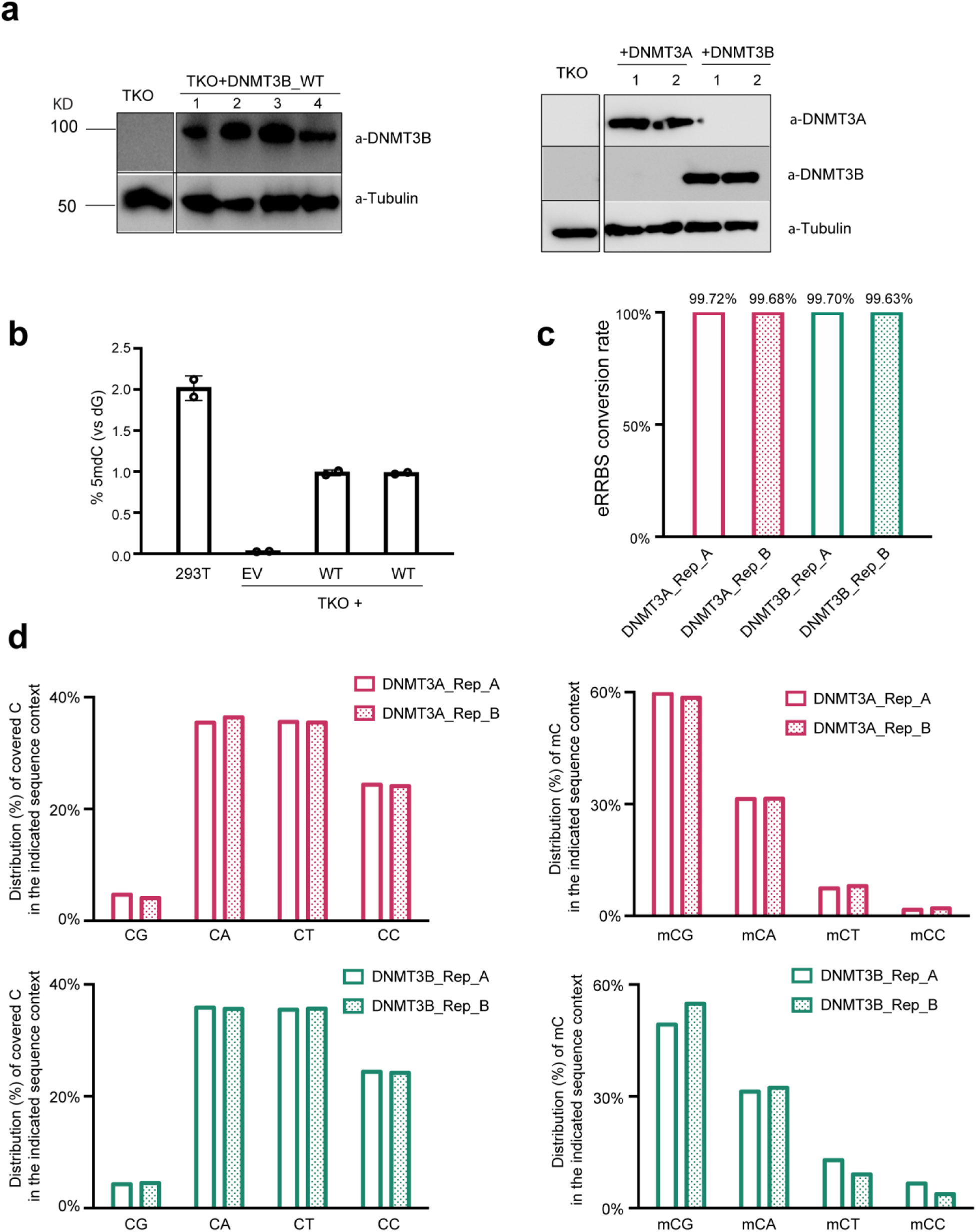
Additional data regarding eRRBS methylation profiles of DNMT3B and DNMT3A. **(a)** Immunoblotting of hDNMT3B and hDNMT3A post-transduction into TKO ES cells reveals their similar expression levels in multiple stable cell lines used for follow-up studies. TKO cells serve as negative. EV, empty vector. **(b)** LC–MS analysis reveals the global 5-methyl-2′-deoxycytidine (5-mdC) levels (indicated as 5-mdC/2′-deoxyguanosine on the y-axis) in the TKO ES cells after stable transduction of empty vector or hDNMT3B (n=3 biological replicates). Data are mean ± SD. **(c)** Bisulfite conversion rates observed in the independent eRRBS sample as determined by the unmethylated lambda DNA spike-in control. **(d)** eRRBS-based methylome profiling shows distribution of the total number of mapped C sites (two left panels) and methylated C (two right panels) with the indicated CpN sequence context in each biological replicate (rep A and B) of TKO lines rescued with either DNMT3A (two upper panels) or DNMT3B (two bottom panels).

**Supplementary Figure 5.**
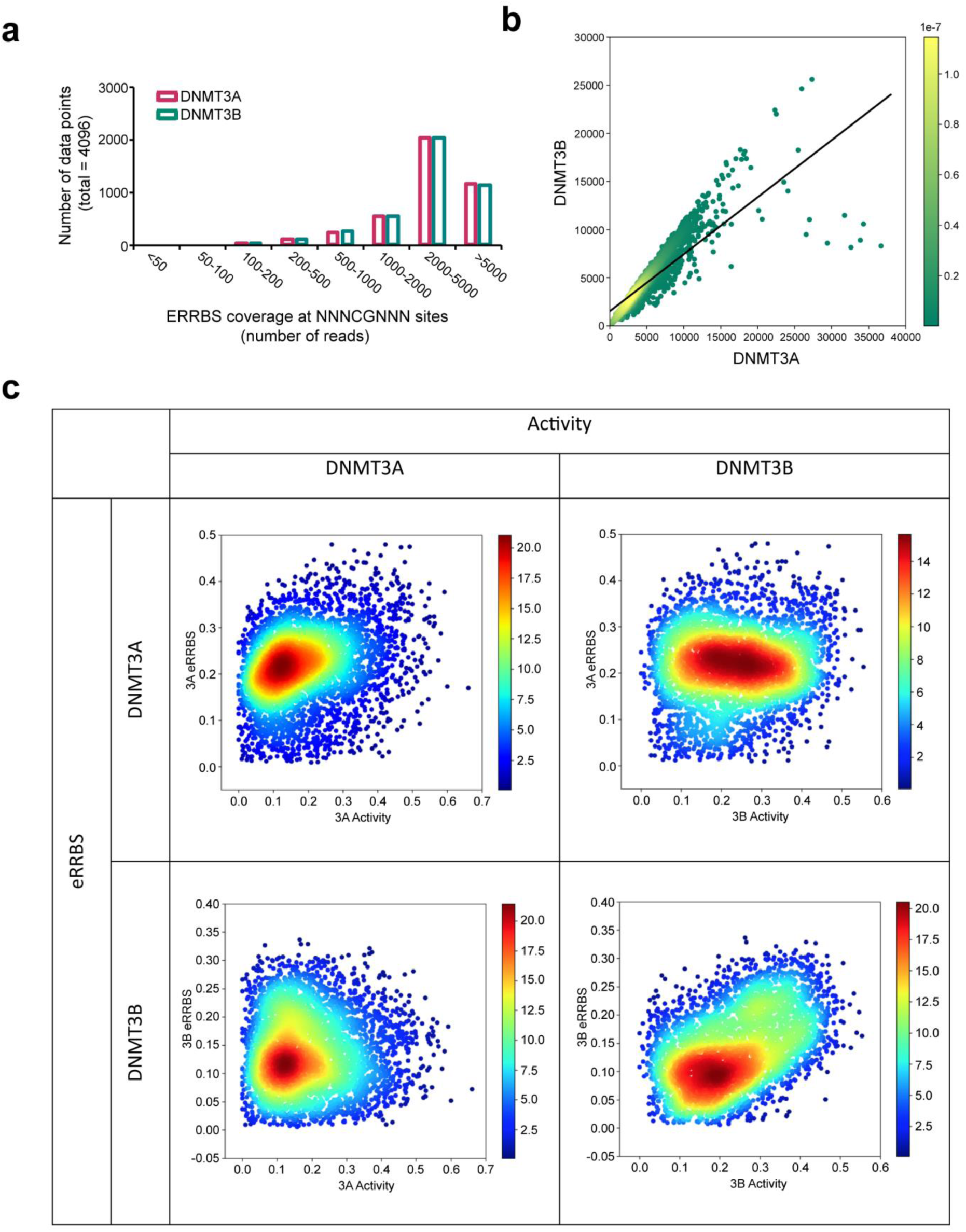
Correlation of the NGS-based activity and eRRBS-based methylation level ratios of DNMT3B/DNMT3A. **(a**,**b)** Bar (b) and Scatter (c) plots showing the coverage of the NNNCGNNN sites in the eRRBS analysis of DNMT3B and DNMT3A. **(c)** Scatter plot showing the correlation between the eRRBS methylation levels at NNNCGNNN sites introduced by DNMT3A and DNMT3B in ESC TKO cells and NGS-based methylation activities of DNMT3A and DNTM3B.

**Supplementary Figure 6.**
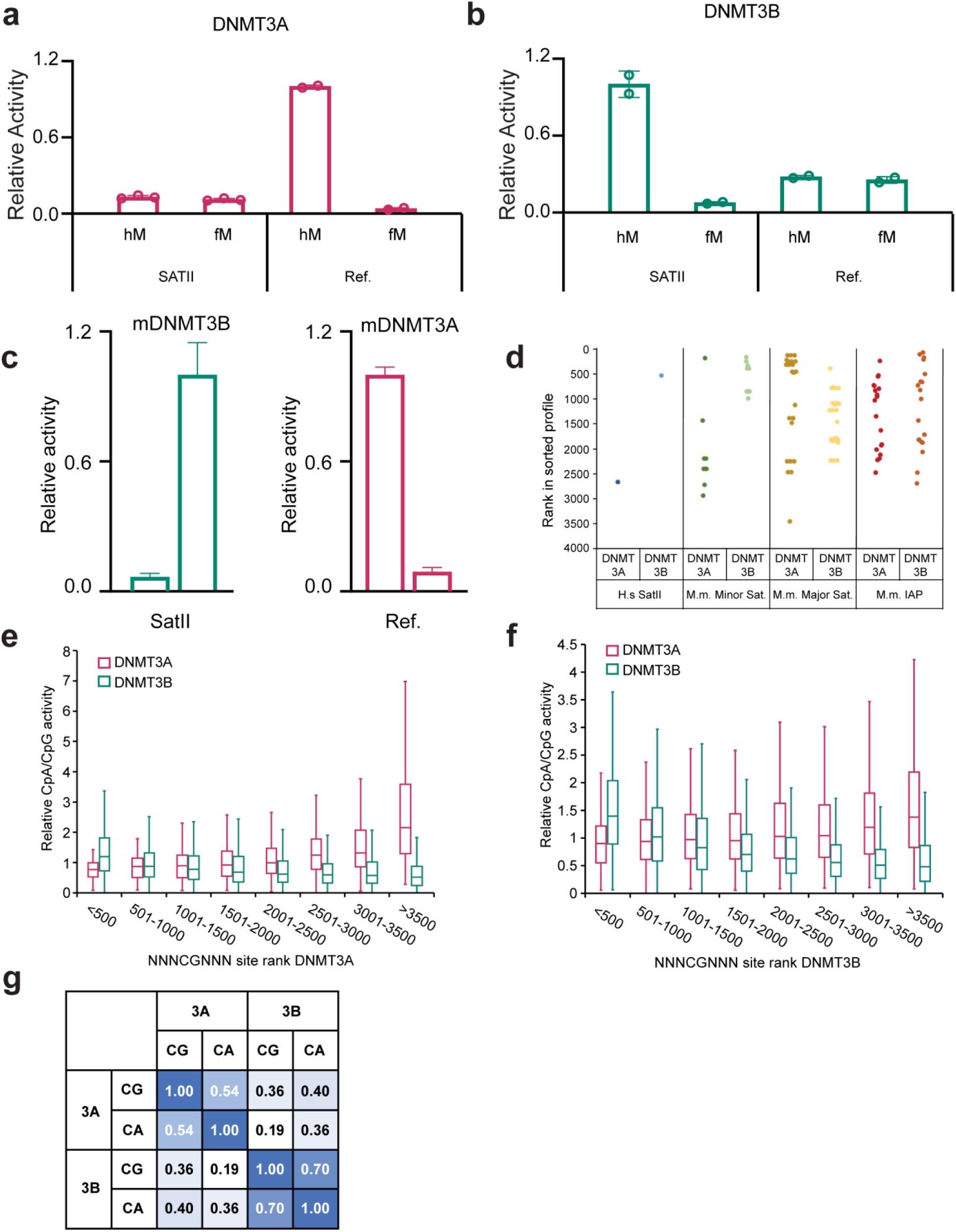
Additional information regarding the flanking sequence preferences of DNMT3A and DNMT3B. **(a**,**b)** Detailed data regarding Fig. 1f. Enzymatic validation of SatII and reference sequences as a substrate for hDNMT3A (a) or hDNMT3B (b), using the MTase domains of DNMT3A and DNMT3B, respectively, analyzed by radioactive methylation assays. Each substrate was used in hemimethylated form and with fully methylated CpG site and the activity observed with the fully methylated CpG site substrate was subtracted from the methylation detected with the hemimethylated form to specifically determine the methylation of the CpG in the upper DNA strand. Average values and standard deviations based on two experimental repetitions are displayed. **(c)** Experimental validation of the SatII preference of mouse DNMT3B (mDNMT3B) by radioactive methylation assays. The methylation activity of mDNMT3B and mouse DNMT3A (mDNMT3A) was normalized to the more active substrate. The figure shows average values and standard deviations based on three independent measurements. **(d)** Sequence preferences of DNMT3A and DNMT3B for CpG sites in human SatII and mouse repetitive elements. All CpG sites from mouse minor satellite repeats (Genbank Z22168.1), major satellite repeats (Genbank EF028077.1) and IAP elements (Genbank AF303453.1) were retrieved. The figure shows the rank of each site in the mDNMT3A and mDNMT3B preference profiles only considering the more preferred DNA strand. A low rank corresponds to high activity. For average values of DNMT3A/DNMT3B ratios refer to Fig. 2d. (**e**,**f**) The relative CpA/CpG methylation activity of DNMT3A or DNMT3B as a function of NGS-based substrate preference rank for DNMT3A (e) or DNMT3B (f). The box indicates the 1^st^ and 3^rd^ quartile with mean indicated. Whiskers show the data range. **(g)** Pearson correlation coefficients of CpG and CpA methylation of mDNMT3A and mDNMT3B in NNCGNN flanking sequence context. Note the higher CpG/CpA correlation in mDNMT3B.

**Supplementary Figure 7.**
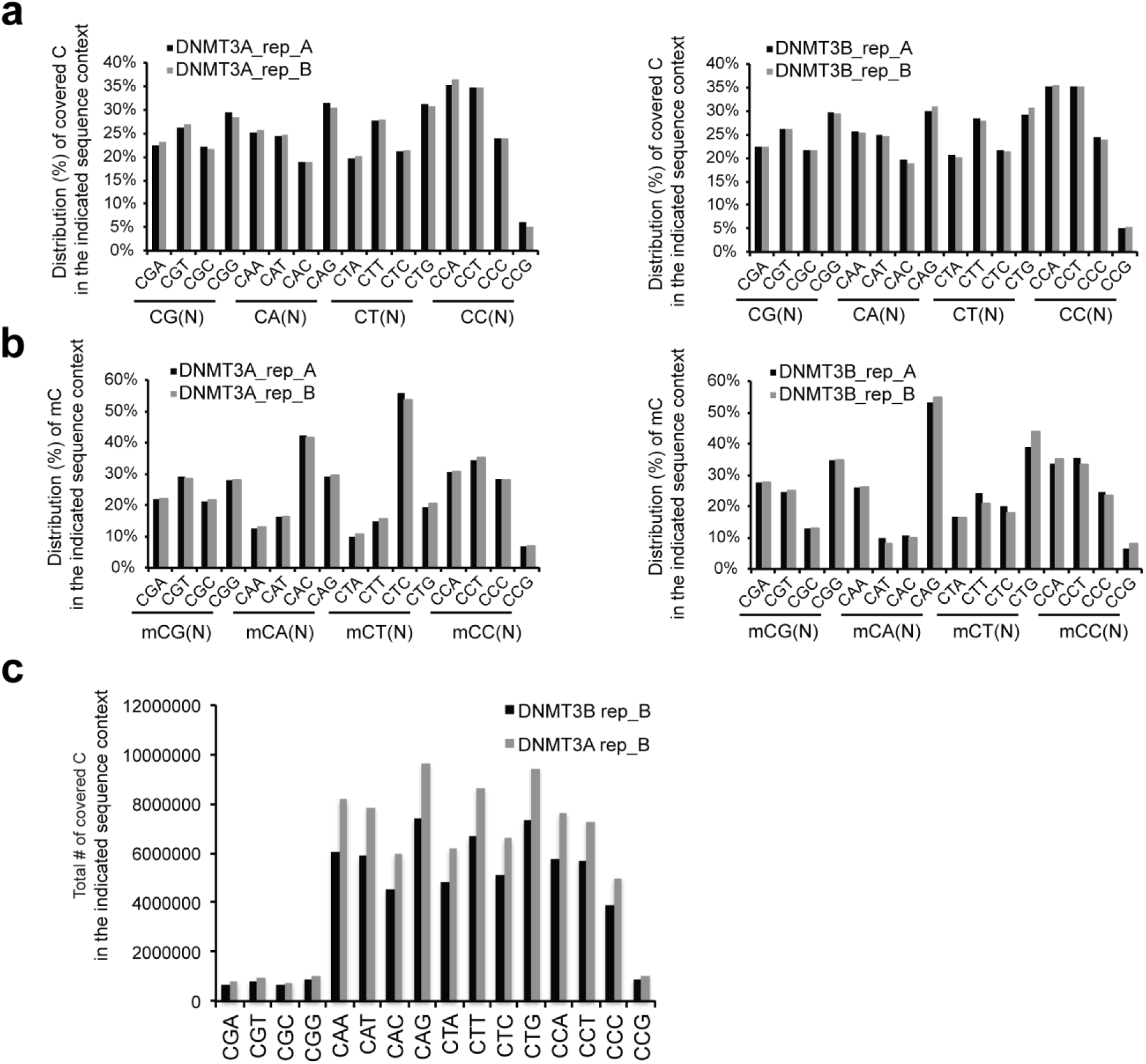
Additional controls regarding the context-dependent DNA methylation by DNMT3A and DNMT3B in cells. **(a**,**b)** eRRBS-based methylome profiling shows distribution of the total number of mapped C sites (panels d) and methylated C (panel e) with the indicated CNN sequence context in each biological replicate (rep A and B) of TKO lines rescued with either WT hDNMT3A (left panel) or hDNMT3B (right panel). **(c)** Total counts for the indicated sequence context as covered by the eRRBS-based methylome profiling in cells.

**Supplementary Figure 8.**
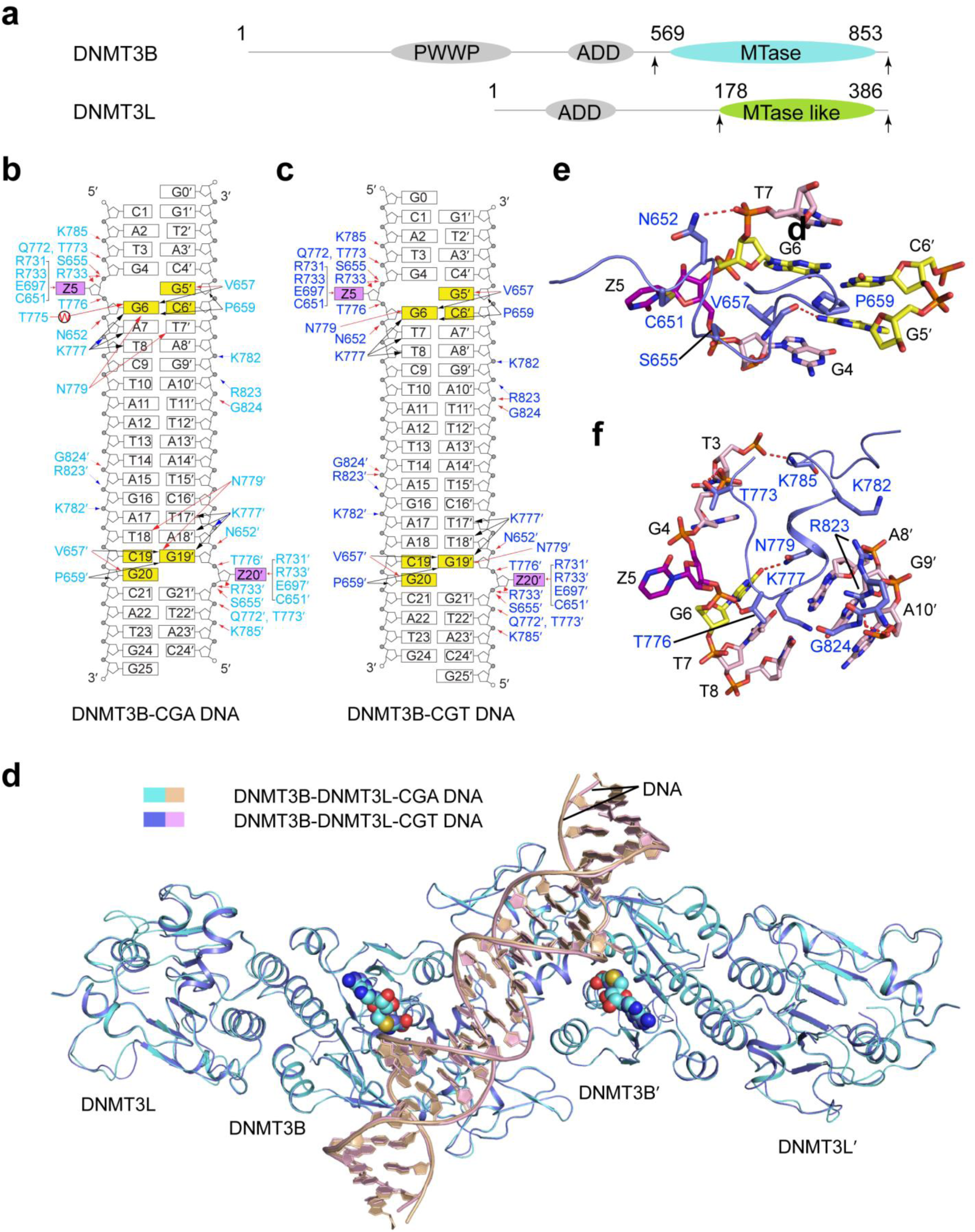
Structural details of the DNMT3B-DNMT3L-DNA complexes. **(a)** Domain architecture of hDNMT3B and hDNMT3L with the C-terminal domains marked with arrowheads. **(b)** Schematic view of the intermolecular interactions between DNMT3B and CGA DNA. The hydrogen-bonding, electrostatic and van der Waals contacts are represented by red, blue and black arrows, respectively. Water-mediated hydrogen bonds are labeled with letter ‘W’. **(c)** Schematic view of the intermolecular interactions between DNMT3B and CGT DNA. The hydrogen-bonding, electrostatic and van der Waals contacts are represented by red, blue and black arrows, respectively. **(d)** Structural overlay of CGA DNA- (cyan) and CGT DNA-bound (slate) DNMT3B-DNMT3L. The bound DNAs are colored in wheat and light pink, respectively. The SAH molecules are shown in spheres, with carbon atoms colored cyan in the DNMT3B-CGA DNA and slate in the DNMT3B-CGT DNA. **(e-f)** Close-up view of the intermolecular interactions between the catalytic loop (**e**), the TRD loop and a loop at the RD interface (**f**) of DNMT3B and DNA. The hydrogen bonds are shown as dashed lines. The ZpG/CpG sites are colored in purple (Z) or yellow.

**Supplementary Figure 9.**
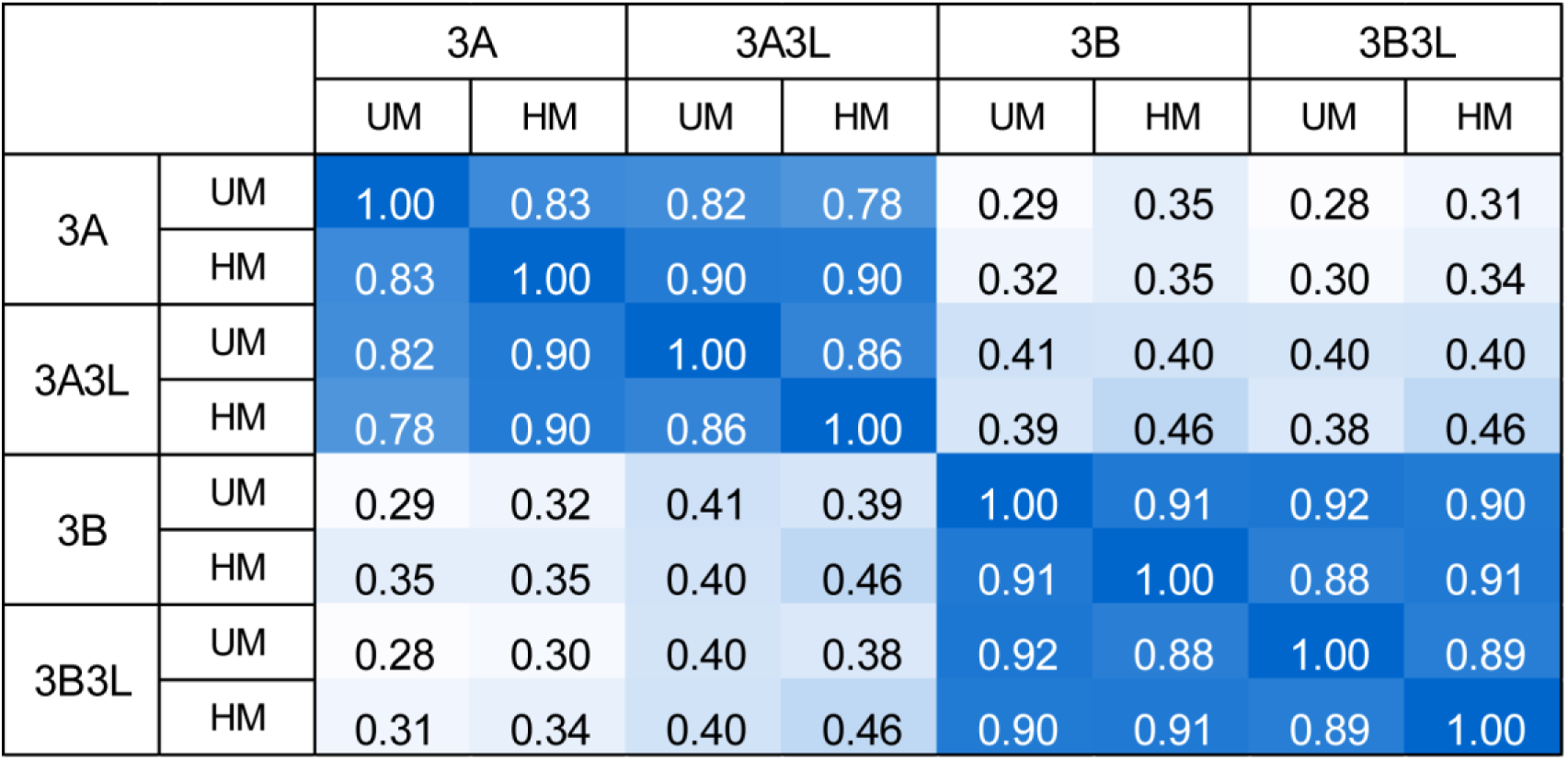
Influence of the methylation level of the CpG site and presence of DNMT3L on the flanking sequence preferences of DNMT3A and DNMT3B. Methylation experiments of random flank substrates with unmethylated (UM) or hemimethylated (HM) CpG site were conducted with murine DNMT3A and DNMT3B in the presence of absence of mDNMT3L. Each experiment was performed in two independent repeats. The figure shows the pairwise Pearson correlation factors of the average methylation levels in each NNCGNN flanking context.

**Supplementary Figure 10.**
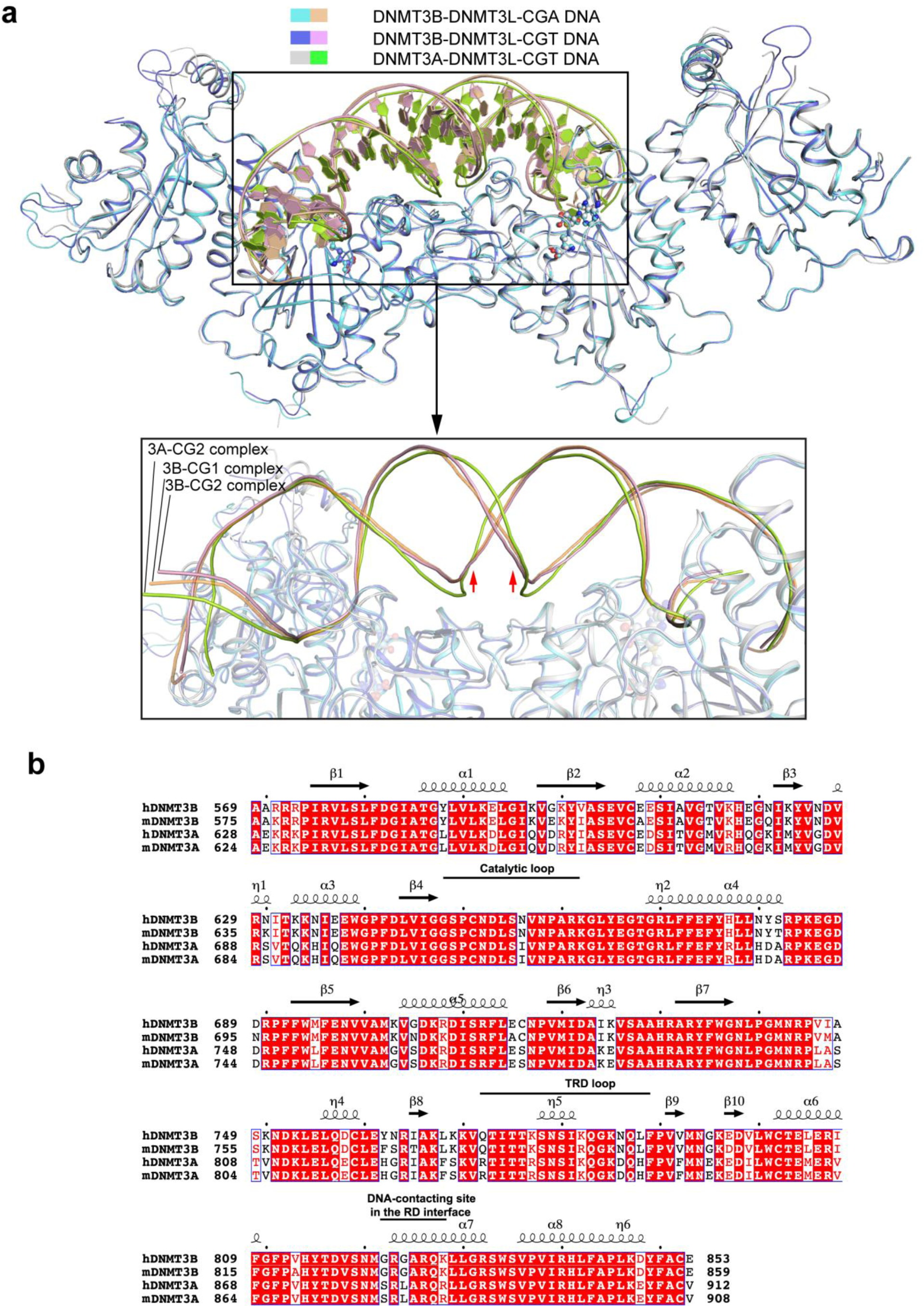
Structural comparison of the DNMT3B-DNMT3L-DNA and DNMT3A-DNMT3L-DNA complexes. **(a)** Structural superposition of the DNMT3B-DNMT3L-CGA DNA, DNMT3B-DNMT3L-CGT DNA and DNMT3A-DNMT3L-CGT DNA. The lower part shows a close-up view of the DNAs bound to DNMT3B or DNMT3A. **(b)** Sequence alignment of DNMT3 proteins from human (hDNMT3B, hDNMT3A) and mouse (mDNMT3B and mDNMT3A). Secondary structures are shown above the aligned sequences. Fully conserved residues are colored in white and highlighted in red shade. Partially conserved residues are colored in red. The residues involved in DNA binding are marked on top of the aligned sequences.

**Supplementary Figure 11.**
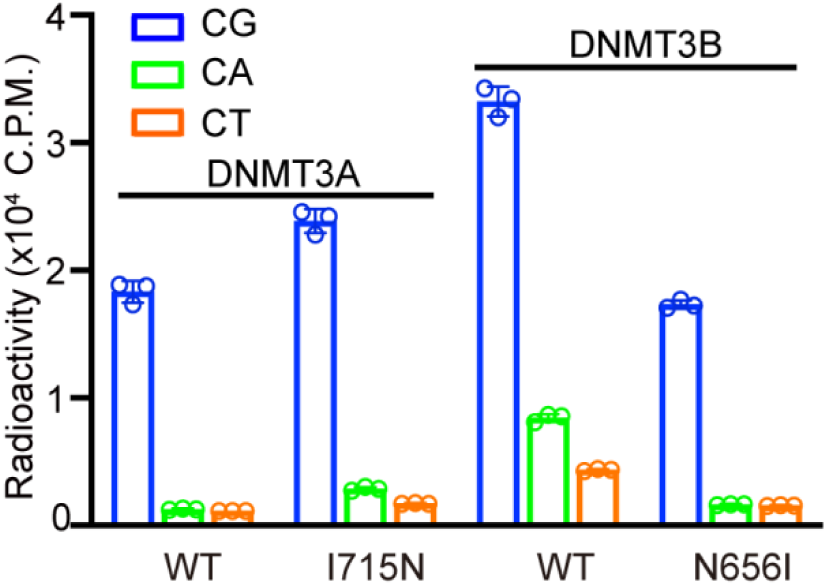
Detailed data regarding Fig. 4e. *In vitro* CpG and CpH methylation of hDNMT3A-hDNMT3L and hDNMT3B-hDNMT3L, WT or mutants on the catalytic loop, were measured using (GAC)_12_, (AAC)_12_ and (TAC)_12_ substrates. The data and error estimate were derived from three independent measurements.

**Supplementary Figure 12.**
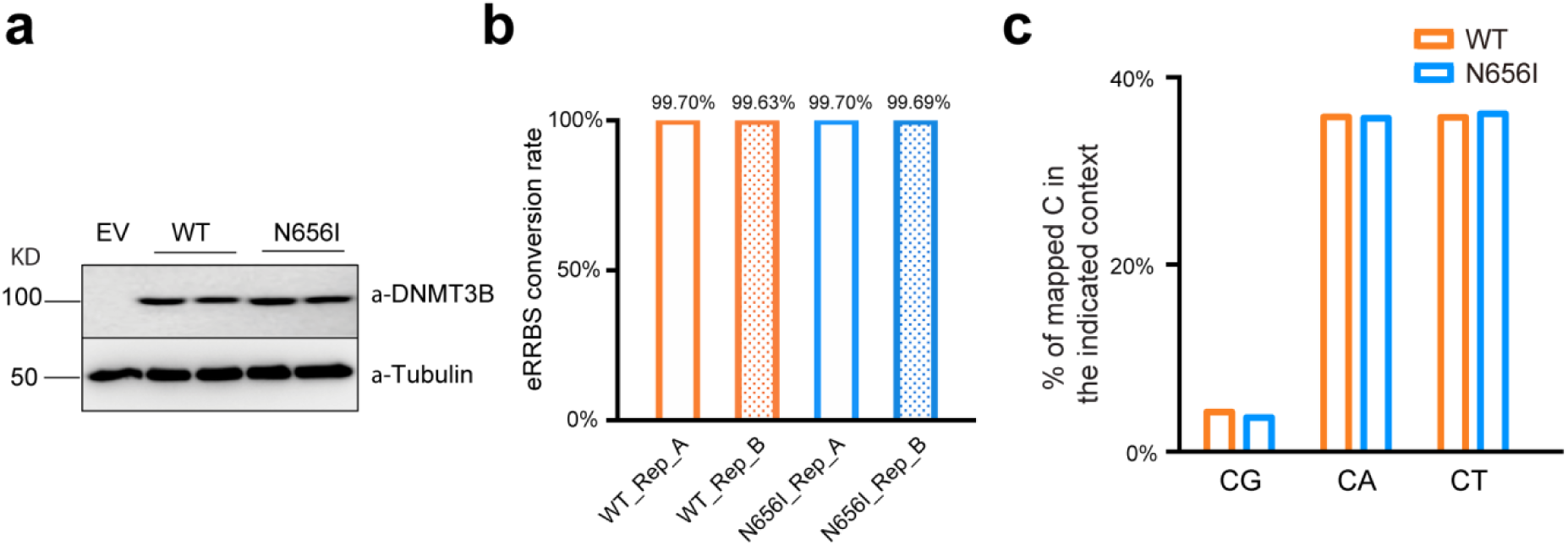
Additional data regarding eRRBS methylation profiles of the DNMT3B N656I mutant. **(a)** Western blot analysis of hDNMT3B, either WT or N656I, post-reconstitution into independently derived TKO lines. **(b)** eRRBS conversion rates of the genomic DNA derived from WT or N656I DNMT3B-transfected TKO cells, as determined by the unmethylated lambda DNA spike-in control. **(c)** eRRBS analysis of WT or N656I hDNMT3B-rescued TKO cells shows similar distribution of the total number of the indicated CpN sites as mapped by sequencing.

**Supplementary Figure 13.**
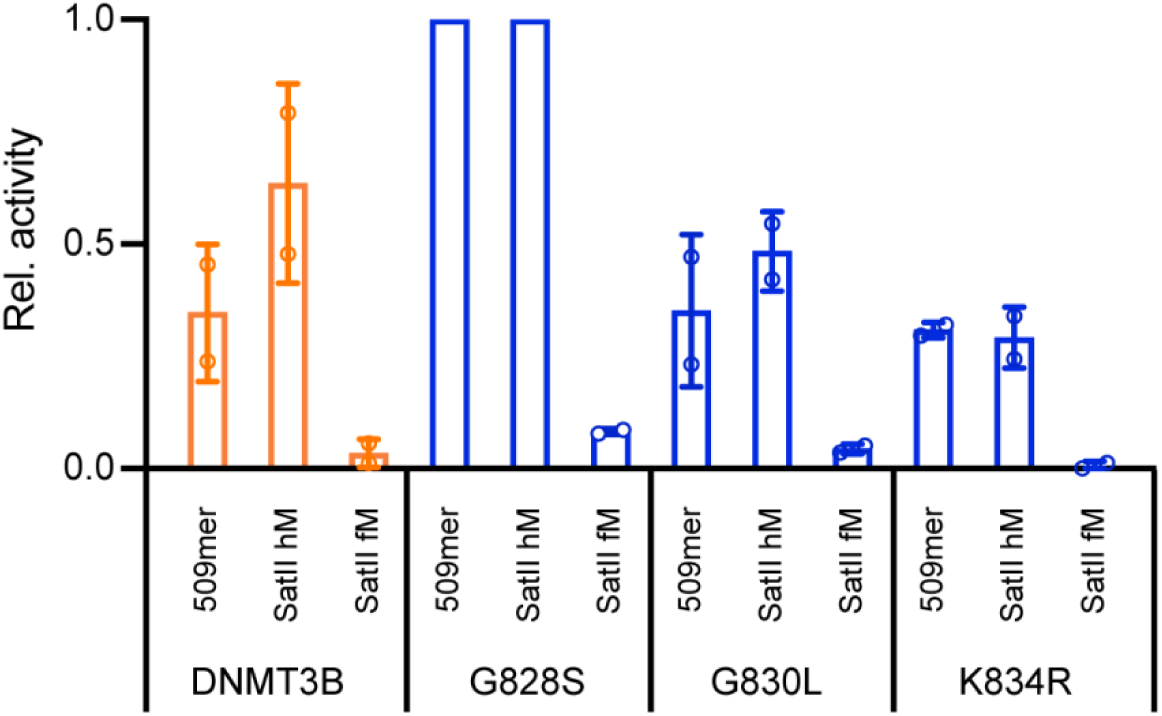
Additional data related to Fig. 4i. The mDNMT3B residues G828, G830 and K834 correspond to hDNMT3B G822, G824 and K828. Methylation activities of DNMT3B mutants were determined on the SatII substrate in hemimethylated (SatII hM) and fully methylated form (SATII fM) and a 500-mer DNA with 58 CpG sites used as “neutral” reference substrate (509mer). The relative preference for the SATII site was calculated as (rate_SATII hM_-rate_SatII fM_)/rate_509mer_ and displayed in relation to WT mDNMT3B (shown in Fig. 4i). The figure shows average values and standard deviations based on two independent measurements.

**Supplementary Figure 14.**
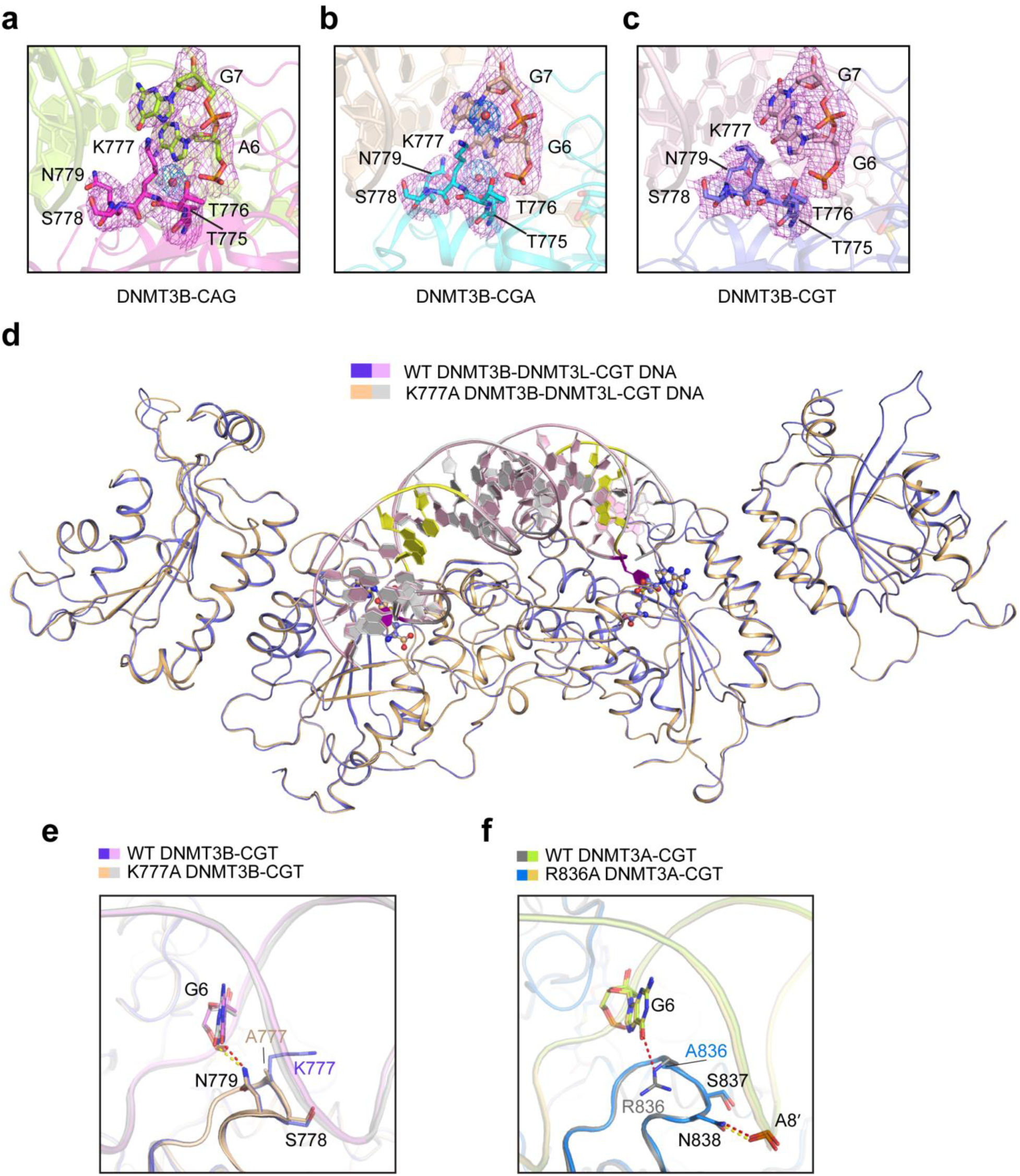
Structural analysis of the TRD loop of DNMT3B in DNA-bound form. **(a-c)** Fo-Fc omit map (violet) for TRD residues T775-N779 of DNMT3B and interacting DNA nucleotides in the DNMT3B-CAG (a), DNMT3B-CGA (b) and DNMT3B-CGT (c) complexes. The Fo-Fc omit map for water molecules (red sphere) are colored blue in (a) and (b). All the omit maps are contoured at 2.0 σ level. **(d)** Structural superposition of the DNMT3B-DNMT3L-CGT DNA and K777A-mutated DNMT3B-DNMT3L-CGT DNA complexes. The CpG/ZpG sites are colored in purple (Zebularine) or yellow. The SAH molecules are shown in sphere representation. **(e)** Close-up view of the aligned WT DNMT3B-CGT and K777A-mutated DNMT3B-CGT complexes, with the hydrogen bonds in the WT and K777A complexes shown as red and wheat dashed lines, respectively. **(f)** Close-up view of the aligned WT DNMT3A-CGT (PDB 5YX2) and R836A-mutated DNMT3A-CGT (PDB 6BRR) complexes, with the hydrogen bonds in the WT and R836A complexes shown as red and lemon dashed lines, respectively.

**Supplementary Figure 15.**
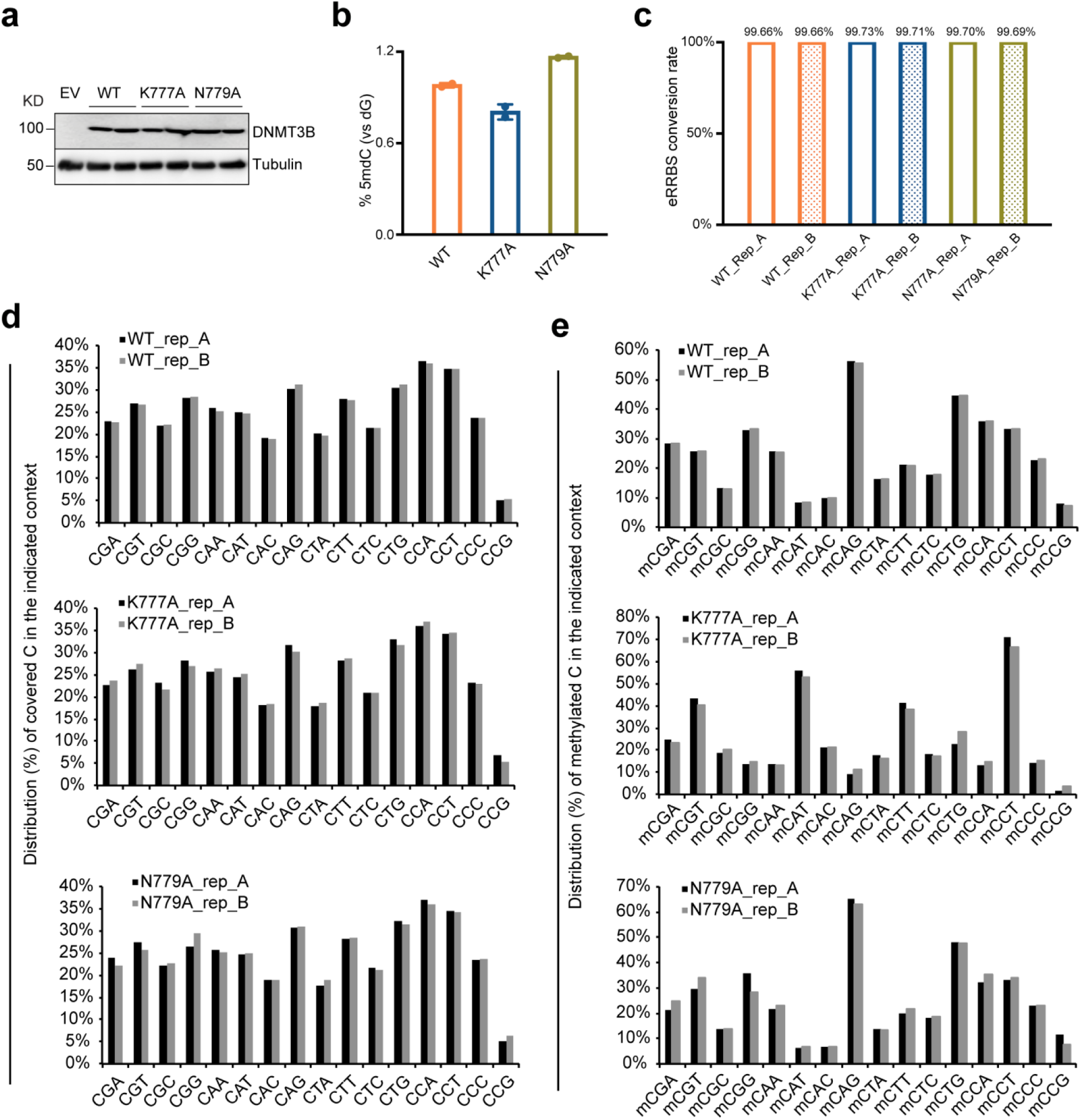
Additional data regarding eRRBS methylation profiles of DNMT3B mutants N777A and K779A. **(a)** Western blot analysis of stably expressed DNMT3B, either WT, K777A or N779A, post-reconstitution into TKO cells. Independently derived lines for each DNMT3B construct were used here and in the following analysis such as eRRBS. **(b)** Liquid chromatography-mass spectrometry (LC-MS) analysis reveals the global 5-mC levels (as calculated by 5-mdC/dG in y-axis) in the TKO ES cells after stable transduction of EV or the indicated DNMT3B (n = 3 biological replicates). Data are mean ± SD. EV, empty vector. **(c)** eRRBS conversion rates of the genomic DNA derived from WT, K777A or N779A DNMT3B-transfected TKO cells, as determined by the unmethylated lambda DNA spike-in control. **(d**,**e)** Bar plots show distribution of the total number of mapped C sites (**d**) and methylated C (**e**) with the indicated CNN sequence context in each biological replicate (rep A and B) of TKO lines rescued with hDNMT3B, either WT (top panel) or the K777A- (middle) or N779A-mutated (bottom).

**Supplementary Figure 16.**
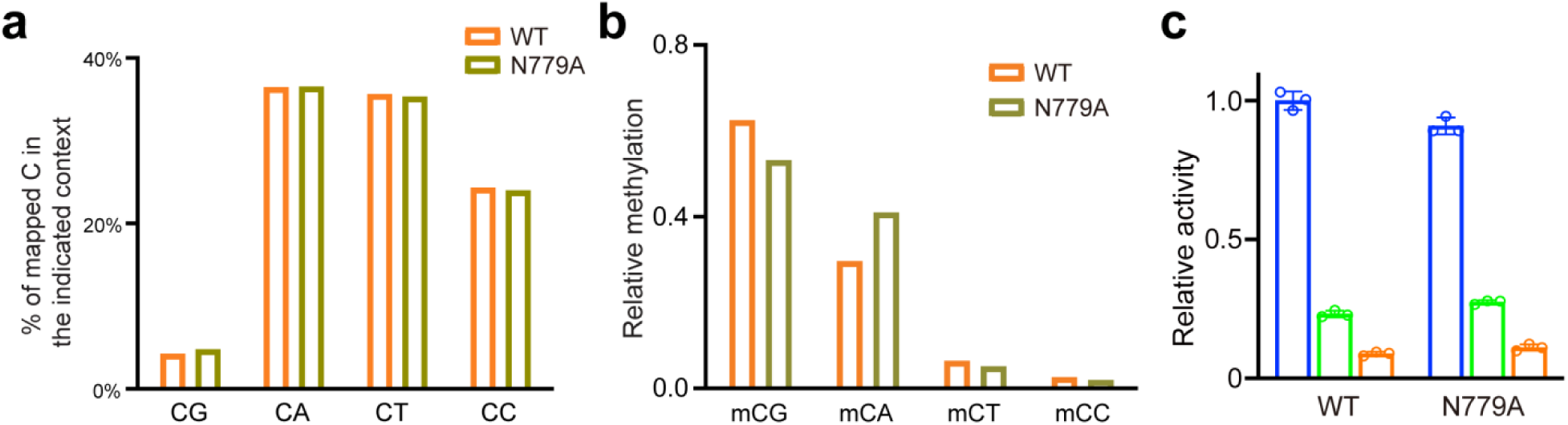
eRRBS and enzymatic analyses of the base preference and selectivity of the DNMT3B N779 mutation. **(a)** eRRBS analysis of WT or N779A hDNMT3B-rescued TKO cells shows similar distribution of the total number of the indicated CpN sites as mapped by sequencing (depth of reads >3 and p < 0.0001). **(b)** eRRBS analysis revealing relative methylation of the indicated context in the TKO cells rescued with WT hDNMT3B or N779A (depth of reads >3 and p < 0.0001). **(c)** *In vitro* CpG and CpH methylation of DNMT3B-DNMT3L, WT or N779A, using (GAC)_12_, (AAC)_12_ and (TAC)_12_ substrates analyzed by radioactive methylation assays.

## Supplementary Tables

**Supplementary Table 1: Overview of the Deep enzymology experiments with DNMT3A and DNMT3B.**

**Supplementary Table 2: Details of the analysis of the deep enzymology experiments. List of NNNCGNNN sites with coverage >50 and very high (>50%) or very low (<1%) methylation by DNMT3A or DNMT3B.**

**Supplementary Table 3: Deep enzymology and eRRBS analyses of mDNMT3A and mDNMT3B in NNNCGNNN sequences.**

**Supplementary Table 4. Summary for the eRRBS-based methylome profiling of TKO cells rescued with WT DNMT3A or DNMT3B, either WT or the N656I mutant.** Two independently derived lines were used for each rescue group.

**Supplementary Table 5. Summary for the eRRBS-based methylome profiling of TKO cells rescued with DNMT3B, either WT, K777A- or N779A-mutated**Two independently derived lines were used for each rescue group. WT DNMT3B-expressing lines used here differ from the two used in Supplementary Table 3.

